# Potassium transporter OsHAK18 mediates shoot-to-root circulation of potassium and sodium and source-to-sink translocation of soluble sugar in rice

**DOI:** 10.1101/2023.03.27.534432

**Authors:** Lirun Peng, Huojun Xiao, Ran Li, Yang Zeng, Mian Gu, Nava Moran, Ling Yu, Guohua Xu

## Abstract

High-affinity potassium (K^+^) transporters HAK/KUP/KT have been identified in all genome-sequenced terrestrial plants. They play important role in K^+^ acquisition, translocation and enhancing salt tolerance. Here we report that the plasma-membrane-located OsHAK18 functions in K^+^ and sodium (Na^+^) circulation and sugar translocation in rice. OsHAK18 is expressed mainly, though not exclusively, in vascular tissues and particularly in the phloem. Knockout (KO) of *OsHAK18* reduced K^+^ concentration in phloem sap and in the root but increased K^+^ accumulation in the shoot of both Nipponbare and Zhonghua11 cultivars, while overexpression (OX) of *OsHAK18* driven by its endogenous promoter increased K^+^ concentration in phloem sap and in roots, and promoted Na^+^ retrieval from shoot to root under salt stress. Split-root experimental analysis of rubidium (Rb^+^) uptake and circulation indicated that *OsHAK18*-OX promoted Rb^+^ translocation from shoot to root. In addition, *OsHAK18*-KO increased while *OsHAK18*-OX reduced soluble sugar content in the shoot and affected oppositely the sugar concentration in the phloem and its content in the root. Moreover, *OsHAK18*-OX increased dramatically grain yield and physiological K^+^ utilization efficiency. Taken together, our results suggest that – unlike other OsHAKs analyzed heretofore – OsHAK18 is critical to K^+^ and Na^+^ re-circulation from shoot to root and enhances the source-to-sink translocation of a photo-assimilate.

**One sentence summary:** Rice potassium transporter OsHAK18 mediates shoot-to-root circulation of potassium and sodium and source-to-sink translocation of soluble sugar which improves potassium use efficiency and grain yield.

## INTRODUCTION

The vascular system of higher plants connects all organs together and carries out long-distance delivery of materials by two parallel structures, xylem and phloem. Xylem conducts water and solutes from root to shoot, while phloem transports mainly photo-assimilates and amino nitrogen (N) compounds in addition to water and minerals from source to sink (White and Ding, 2022). K^+^ is the most abundant cation in the sap of xylem or phloem.

Potassium (K^+^) is the most abundant cation in plants in general, and is critical in maintaining ion charge balance, regulating turgor pressure and enzymes activity, as well as in promoting the mobilization of other nutrients and sugar (Anschutz et al., 2014; Li et al., 2018). K^+^ is a highly mobile cation, it circulates in plants; regulating photosynthetic carbon translocation from source to sink. About 40-90% of K^+^ transported from the root via xylem to aerial parts is circulated from the shoot to roots along with photosynthates, sugars and other nutrients (Lu et al., 2005; Peuke et al., 2011). Such circulation may be an important feedback signal regulating root K^+^ uptake and smoothing out the fluctuations of external K^+^ supply (Marschner et al., 1996). K^+^ gradients serve as a mobile energy source in plant vascular tissues and K^+^ circulating in the phloem transports energy stored in the K^+^ gradient between phloem cytosol and the apoplast (Gajdanowicz et al., 2010; Dreyer et al., 2017). K^+^ and sodium (Na^+^) ions share great similarity in physical and chemical characteristics, thus, enhancing K^+^ absorption can impair Na^+^ accumulation which is one of the key strategies for salt stress tolerance in glycophytes (Maathuis and Amtmann, 1999; Munns and Tester, 2008; Shabala and Cuin, 2008).

K^+^ is absorbed by roots from the soil and circulated within the plant by multiple families of K^+^ channels and transporters. The Shaker family of voltage-gated K^+^ channels and high-affinity K^+^ transporters (HAK/KUP/KTs) are major proteins underlying intercellular K^+^ transport. K^+^ entry into roots when the external K^+^ concentration at the root surface is high, occurs via the K^+^ channel AKT1 down the K^+^ electrochemical potential, while at a low external K^+^ concentration, a high-affinity K^+^ uptake occurs via HAK/KUP/KT family members such as via AtHAK5 in Arabidopsis (Rubio et al., 2010) and OsHAK1 and OsHAK5 in rice (Yang et al., 2014; Chen et al., 2015). At the root endoderm, the hydrophobic Casparian strip prevents apoplastic movement of K^+^ into xylem sap, therefore, K^+^ must enter the symplast of the endodermis and then be released into xylem sap. Protoplasts of stellar cells (the root-xylem-enwrapping parenchyma cells) possess voltage-gated <u>S</u>haker-like <u>K</u>^+^ <u>o</u>utwardly rectifying channels (SKOR) (Wegner and Raschke, 1994; Roberts and Tester, 1995). These SKOR channels subserve K^+^ release into the xylem sap toward the shoots (Gaymard et al., 1998). Moreover, high-affinity K^+^ transporters like AtKUP7 and OsHAK5 which are also expressed in stellar cells, contribute significantly to the root-to-shoot K^+^ translocation via xylem, particularly when K^+^ supply is low (Yang et al., 2014; Han et al., 2016).

Phloem is comprised of longitudinally aligned sieve elements (SEs) and adjoining companion cells (CCs). It is divided into three functional domains, collection phloem, transport phloem and unloading phloem (De Schepper et al., 2013). Loading photo-assimilates and inorganic ions, mainly sucrose, into SE/CC complexes of the collection phloem at source leaves and releasing them from the unloading phloem in sink organs generate an osmotically derived hydrostatic pressure gradient (Turgeon, 2010). This hydrostatic pressure gradient is considered to be the major driving force for the phloem bulk flow in herbaceous plants (Crafts and Crisp, 1972; Knoblauch and Peters, 2016). High K^+^ concentration in sieve tubes of phloem and a steep transmembrane pH gradient are required for sucrose/H^+^ co-transporter (SUT1) to load apoplastic sucrose into the sieve tubes at source leaves (Ayre, 2011; Julius et al., 2017). However, even with impaired SUT1, the bulk flow in phloem was preserved due to the elevation of K^+^ concentration at the source which served as the main osmoticum compensating for decreased loading of sucrose (Babst et al., 2022).

The transport phloem connecting the collection phloem and the unloading phloem allows a “leak out” of solutes which partially nourish the tissues in the plant axis (van Bel, 2003), or are reloaded back into the tube (Pate and Dieter Jeschke, 1995; Marschner et al., 1996). The leakage of K^+^ from the companion cells of the transport phloem is conducted by the AKT2 channel of the Shaker family (Deeken et al., 2002) shown, by genetic manipulation and computational modeling, to function here as a K^+^-release channel (Gajdanowicz et al., 2010). The switch of the AKT2 gating from voltage-rectifying to non-rectifying, which allows K^+^ release, occurs by channel phosphorylation (Michard et al., 2005).

Solutes including K^+^ are loaded into the collecting phloem via both symplastic and apoplastic routes. In the symplastic route, plasmodesmata from mesophyll cells allow the passage of solutes directly into the companion cells, while transporters are required in the apoplastic route. Whether K^+^ loading into the collecting phloem involves a K^+^ channel is uncertain, though a phloem-expressed K^+^ channel in maize (KZM1) can mediate K^+^ influx current in *Xenopus* oocytes (Philippar et al., 2003). Computational modeling demonstrated that energy-dependent K^+^ transport is required for K^+^ loading into phloem in source leaves (Turgeon and Ayre, 2005). However, the identity of K^+^ transporter(s) involved in phloem K^+^ re-circulation in plants is not known.

In this study, we provide molecular, physiological and genetic evidence that OsHAK18 (accession no. AK065464), a member of cluster III of HAK/KUP/KT family (Supplemental Figure S1), plays a major role in the re-cycling of K^+^ and Na^+^ from shoot to root in rice. OsHAK18 is expressed abundantly in rice mesophyll cells and vascular tissues, functions in efficient utilization of K^+^ and limiting Na^+^ accumulation in the shoot, and contributes to source-to-sink translocation of K^+^-accompanied soluble sugar used in the production of grains.

## RESULTS

### Tissue and cellular localization of *OsHAK18* expression in rice

Plant HAK transporters have been shown to contribute to root K^+^ uptake and translocation of K^+^ from root to shoot under varied supply of [K^+^]. By checking the expression pattern of rice HAK transporter genes via publicly released microarray data (RiceXPro, http://ricexpro.dna.affrc.go.jp/), we found that *OsHAK18*, unlike other HAK family members, is minimally transcribed in the root, suggesting that its function may divert from what is currently known about this family. Before exploring the physiological function of this gene, we first examined the organ-specific expression pattern of this gene by qRT-PCR in 3-week-old seedlings. Indeed, the transcript level is low in the root and leaf sheath but high in the leaf blade (Supplemental Figure S2, A-C).

To examine further the tissue and cellular localization of this gene in rice, the GUS (β-glucuronidase) reporter gene was fused to the putative promoter of *OsHAK18* (2000 bp upstream of the translational start codon), and then transgenic rice plants harboring the *ProOsHAK18:GUS* construct were generated. Histochemical analysis of these transgenic rice plants showed that *OsHAK18* is expressed in the vascular cylinder of roots, basal nodes and leaves, as well as in mesophyll cells (MCs; Supplemental Figure S2D, a-c; Figure 1). OsHAK18 is also expressed in the glumous flower (Supplemental Figure S2Dd).

**Figure 1.**
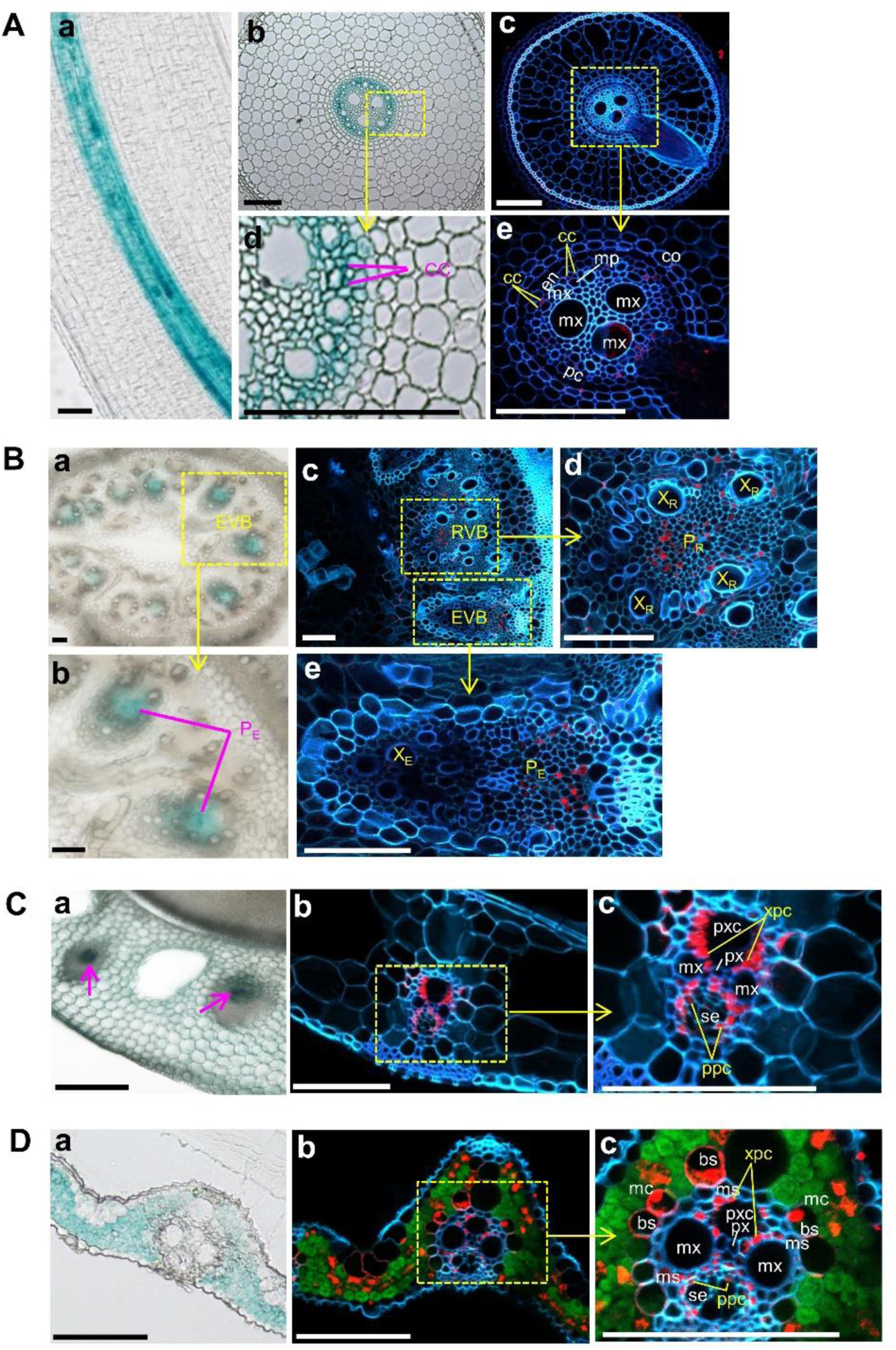
Tissue and cellular localization of *OsHAK18* in rice. The localization is based on the detection of GUS reporter staining in transgenic rice plants harboring the *ProOsHAK18:GUS* construct and by *OsHAK18* immunostaining. A, crown root. B, basal node. C, leaf sheath. D, leaf blade. The GUS staining is seen in the transmitted-light images, Aabd, Bab, Ca, Da. The immunostaining - in confocal fluorescence images, Ace, Bcde, Cbc, Dbc. In the immunostained cross sections, the red, green and blue colors represent the expression of *OsHAK18*, chlorophyll autofluorescence and cell wall autofluorescence, respectively. bs, vascular bundle sheath; cc, phloem companion cell; co, cortex; en, endodermis; EVB, enlarged vascular bundle; mc, mesophyll cell; mp, metaphloem; ms, mestome sheath; mx, metaxylem vessel; pc, pericycle; P_E_, phloem of EVB; P_R_, phloem of RVB; px, protoxylem; pxc, protoxylem cavity; RVB, regular vascular bundle; se, sieve element; ppc, phloem parenchyma cell; X_E_, xylem of EVB; xpc, xylem parenchyma cell; X_R_, xylem of RVB. Scale bars = 100 μm.

To further examine the expression of *OsHAK18* at a cellular level, an immunostaining analysis was performed. In both roots and basal nodes, *OsHAK18* expression was restricted to the phloem. Specifically, *OsHAK18* was expressed in the companion cells (CCs) of the root phloem (Figure 1A, d and e) as well as in the phloem of the regular vascular bundle (RVB) and enlarged vascular bundle (EVB; Figure 1B, c-e). Unlike that found in roots and basal nodes, *OsHAK18* was expressed in both the phloem parenchyma cells (PPCs) and xylem parenchyma cells (XPCs) of leaf sheaths and leaf blades (Figure 1, Cc and Dc). Additionally, in leaf blades, it was also transcribed in vascular bundle sheath (BS) layer and MCs (Figure 1Dc).

### OsHAK18 is located in the rice protoplast plasma membrane and mediates K^+^ and Na^+^ import in yeast

To determine the subcellular localization of OsHAK18, we transiently co-expressed in rice protoplasts *OsHAK18* fused to *eGFP* at its N- or C-terminal, and *OsRac1* fused to mCherry as the plasma membrane marker (Figure 2). eGFP alone, the product of an ‘empty’ vector, diffused within the cytosol, while the green signal of the eGFP-fused OsHAK18 overlapped very well with the plasma membrane marker, irrespective of the OsHAK18 terminus to which eGFP was fused (Figures 2, A and B).

**Figure 2.**
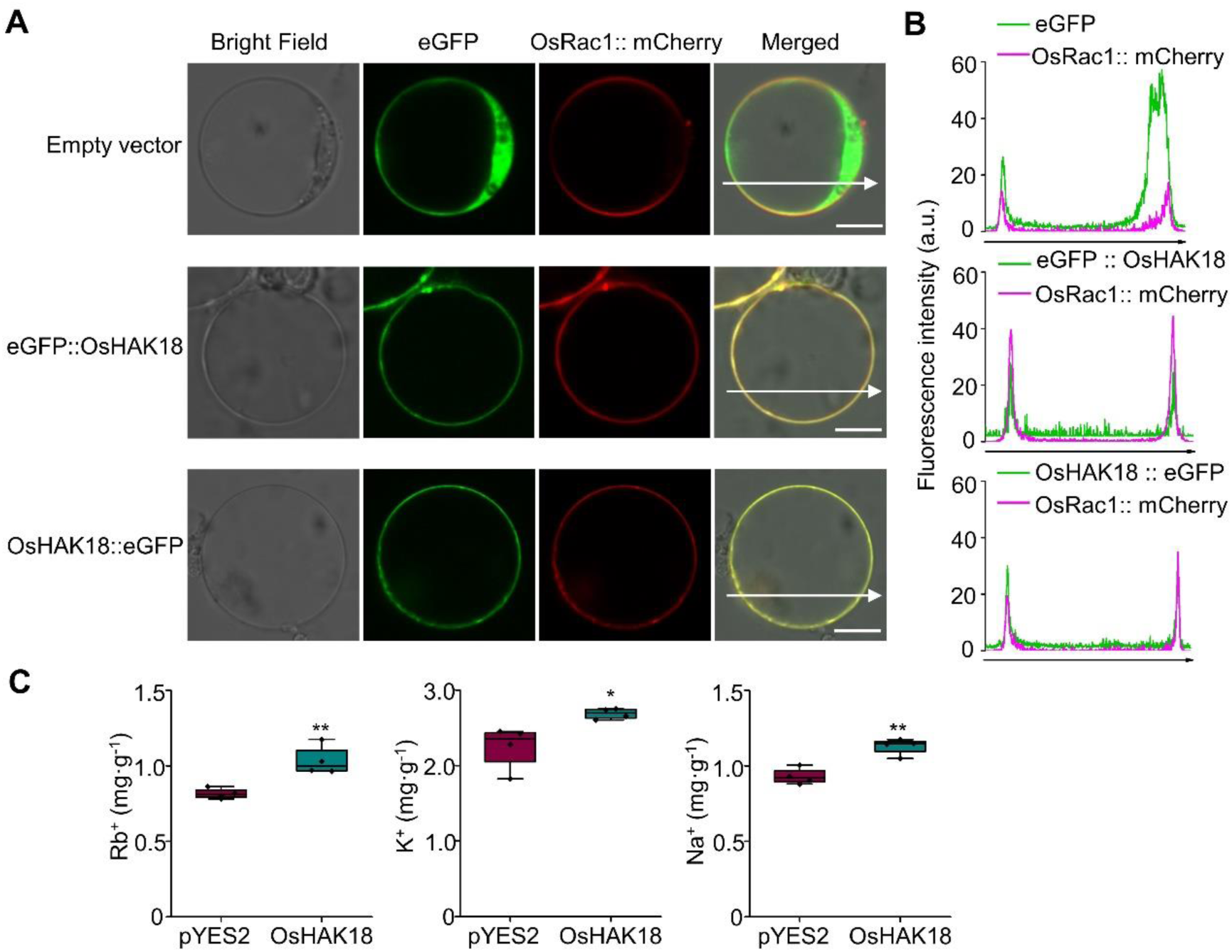
Subcellular localization of OsHAK18 and evaluation of the K^+^ transport activity of OsHAK18 in yeast. A, An expression of eGFP::OsHAK18 or OsHAK18::eGFP fusion proteins in rice protoplasts. OsHAK18 was co-localized with the plasma membrane-localized GTPase OsRac1. Scale bars = 10 μm. B, Line scans of rice protoplasts co-expressing OsHAK18– eGFP and OsRac1. C, Cells of G19 transformed with the empty vector pYES2 or OsHAK18 were cultured in liquid SC-U medium supplemented with RbCl for 12 h, KCl or NaCl for 24 h. In the box plots, the symbols represent individual data, whiskers represent minimum and maximum data values, the boxes are delimited by the first and the third quartiles, and the horizontal line within the box indicates the median. Significant differences from pYES2 are indicated by asterisks (*, P<0.05; **, P<0.01). Non-paired, two-tailed, equal-variance t-test was used for the statistical analysis.

To test whether OsHAK18 can mediate K^+^ uptake, we used the yeast strain R5421, the *trk1 trk2* double mutant defective in high-affinity K^+^ uptake, for the functional complementation assay of the transporter. Both OsHAK18 and OsHAK5 (as a positive control of a high-affinity K^+^ transporter) were cloned into three different yeast expression vectors, p416GPD, pDR196 and pYES2, and transformed into R5421. As expected, OsHAK5 could rescue the growth defect of R5421 mutant at low external K^+^ medium (0.1 mM and 1 mM), however, OsHAK18 did not complement the defective growth of R5421 (Supplemental Figure S3), implying that OsHAK18 cannot serve as a high affinity K^+^ transporter, at least in yeast.

*OsHAK18* was also expressed in a Na^+^-sensitive yeast strain G19. OsHAK18 increased by about 25% the Rb^+^ content of G19 supplied with Rb^+^ (Figure 2C). Moreover, OsHAK18 could elevate K^+^ and Na^+^ content of G19 supplied with KCl and NaCl, respectively (Figure 2C). These results suggest that OsHAK18 is able to transport both Na^+^ and K^+^.

### OsHAK18 affects significantly K^+^ distribution between root and shoot

The GUS expression profile, subcellular location and the Rb^+^ transport activity of OsHAK18 in G19 yeast suggest that OsHAK18 mediates K^+^ translocation in rice. To delve into the physiological function of this transporter, we obtained two different *Tos17* insertion mutants of *OsHAK18*-knockout (KO) in genetic background of cv. Nipponbare from NIAS institute in Japan and cv. Zhonghua11 from Huazhong Agricultural University in China, and verified their *Tos17* insertional sites (Supplemental Figure S4, A-C). In addition, we generated *OsHAK18* self-promoter-controlled overexpression lines (OX) in cv. Nipponbare (Supplemental Figure S4, D-F).

K^+^ concentration [K^+^] has been determined in all genotypes, in the root and in two parts of the shoot, the leaf sheath (L.S.) and the leaf blade (L.B.) at two external K^+^ concentrations ([K^+^]_EXT_) outside the root (1 mM or 0.1 mM). In the wild type (WT) plants, the L.S.’s [K^+^] was about 30 mg K^+^ per g DW, roughly 20-100% higher than the L.B.’s [K^+^] (Figure 3, A-D). In contrast, in the *OsHAK18*-KO plants, at the normal [K^+^]_EXT_ of 1 mM, the root’s [K^+^] was reduced relative to WT’s, but [K^+^] in the KO’s shoot did not differ from WT’s shoot (Figure 3A). At [K^+^]_EXT_ of 0.1 mM, [K^+^] in the KO’s root was also reduced relative to WT’s, but in contrast, in the KO’s leaf sheath, [K^+^] was elevated appreciably compared to WT’s (Figure 3C).

**Figure 3.**
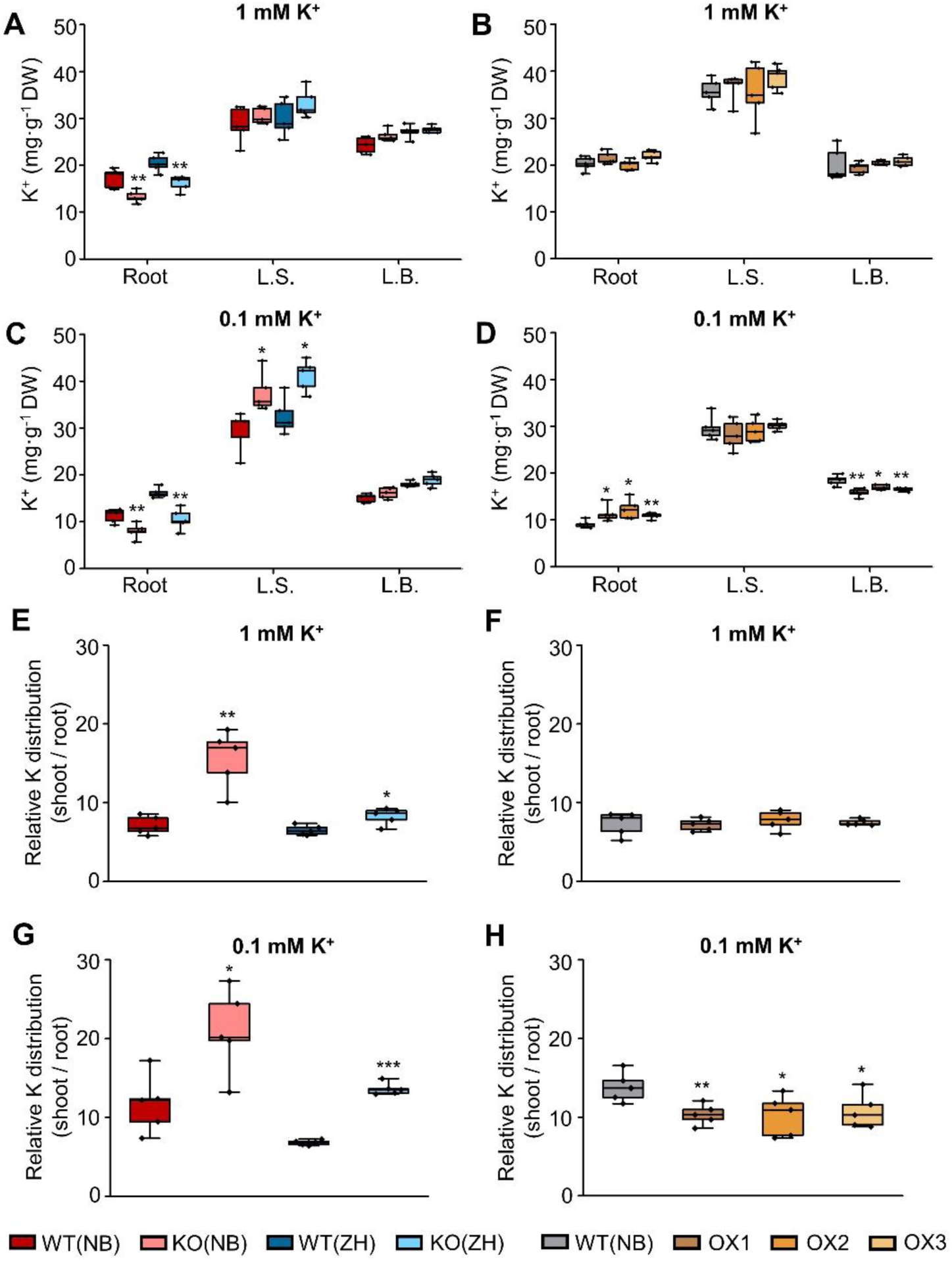
Effect of *OsHAK18* expression level on K^+^ distribution at the seedling stage. Ten-day-old seedlings were grown in IRRI solution containing 1 mM K^+^ for 4 weeks, then transferred to 1 mM K^+^ (A, B, E, F) or to 0.1 mM K^+^ (C, D, G, H) solutions for 2 weeks. A-D, K^+^ concentration in root, leaf sheath (L.S.) and leaf blade (L.B.). E-H, Relative K^+^ distribution (shoot / root ratio of total K^+^ content). Detailed data on total K^+^ content in root and shoot are provided in Supplemental Figure S6. The box plots and t-tests are as in Figure 2. Significant differences from the corresponding wild type (WT) are indicated by asterisks (*, P<0.05; **, P<0.01; ***, P<0.001).

The biomass of the OsHAK18-KO plants was smaller than that of WT (Supplemental Figure S5, A, B, E and F), such that the total K^+^ content (biomass* [K^+^]) in any of the separate parts of the KO plant – root, L.B., L.S. – was appreciably smaller than in the WT, and by a larger percentage in the root (75 %) than in the shoot (50%; Supplemental Figure S6A). A similar decrease (in %) in the total K^+^ content was observed in KO vs the WT plants at the low [K^+^]_EXT_ of 0.1 mM, even though at this low [K^+^]_EXT_ the total K^+^ content was lower in both genotypes, relative to the regular [K^+^]_EXT_ (Supplemental Figure S6B). Therefore, the shoot-to-root ratio of the total K^+^ content (a measure of total K^+^ distribution in the plant) increased appreciably, also irrespective of [K^+^]_EXT_ (Figure 3, E and G). In summary, compared to WT, *OsHAK18*-KO dramatically increased the [K^+^] in the shoot at [K^+^]_EXT_ of 0.1 mM (Figure 3C) and shifted the relative distribution of K^+^ to the shoot, irrespective of the [K^+^]_EXT_ (Figure 3, E and G; see Discussion).

The OsHAK18 function was also analyzed in three representative OX lines. The biomass of all parts of the OX plants was also smaller than WT’s (Supplemental Figure S5, C, D, G and H). While at [K^+^]_EXT_ of 1 mM, OX did not affect [K^+^] neither in the root nor in the shoot (Figure 3B), at the low [K^+^]_EXT_ of 0.1 mM, OX increased [K^+^] in the root and decreased it in the leaf blade (Figure 3D). Thus, with 1 mM K^+^ in the growth solution, OX decreased the total K^+^ content both in the shoot and the root (Supplemental Figure S6C), but it did not affect the relative distribution of total K^+^ between the shoot and the root relative to WT’s (Figure 3F). In contrast, at [K^+^]_EXT_ of 0.1 mM, OX diminished the total K^+^ content only in the shoot (Supplemental Figure S6D) and thus decreased the shoot-to-root ratio of the total K^+^ content (Figure 3H). These results suggest that OsHAK18 affects appreciably the distribution of K^+^ between the root and the shoot, particularly under a limited K^+^ supply.

### OsHAK18 does not contribute to root K^+^ uptake and transport from root to shoot

In the root, *OsHAK18* is expressed only in the stele, not in epidermal and cortex cells (Figure 1A, a-c), which suggested that OsHAK18 may not contribute to K^+^ uptake. To test this, we determined the short-term K^+^ acquisition rate (normalized to the root weight) at 1 mM or 0.1 mM [K^+^] in the growth medium, which was determined by K^+^ depletion experiments from the external solution. The K^+^ acquisition rate during a 12-h assay, did not differ among all the rice genotypes, WT, KO and OX at both [K^+^]_EXT_, even if it was roughly three times lower at the low [K^+^]_EXT_ compared to the regular [K^+^]_EXT_ (Supplemental Figure S7). These results exclude a direct role of OsHAK18 in the root K^+^ uptake in rice.

To examine directly the OsHAK18 role in the xylem transport of K^+^, we first analyzed K^+^ concentration in the xylem sap of all our rice genotypes grown in 1 or 0.1mM K^+^. To generate similar starting conditions for both treatments, the roots were immersed for three hours in a medium without K^+^ (because otherwise, they would start with different root [K^+^] as a result of the different growth conditions (Figure 3, A-D), which would generate bias in the xylem sap [K^+^]). We collected the xylem sap initially in one-h intervals for three consecutive hs at nominally zero K^+^ and subsequently – for additional three hs with either 1 or 0.1 mM K^+^ restored to the bath. During the K^+^-deprivation period, the KO plants grown in 1 mM K^+^ had a lower xylem sap [K^+^] than the corresponding WT only during the first h (Figure 4A), while in KO grown in 0.1 mM K^+^, this difference in the xylem sap [K^+^] lasted for twice as long (Figure 4B). However, after 3 hours, the root [K^+^] became the same in both treatment groups (a suitable starting point for assaying the effect of K^+^ resupply).

**Figure 4.**
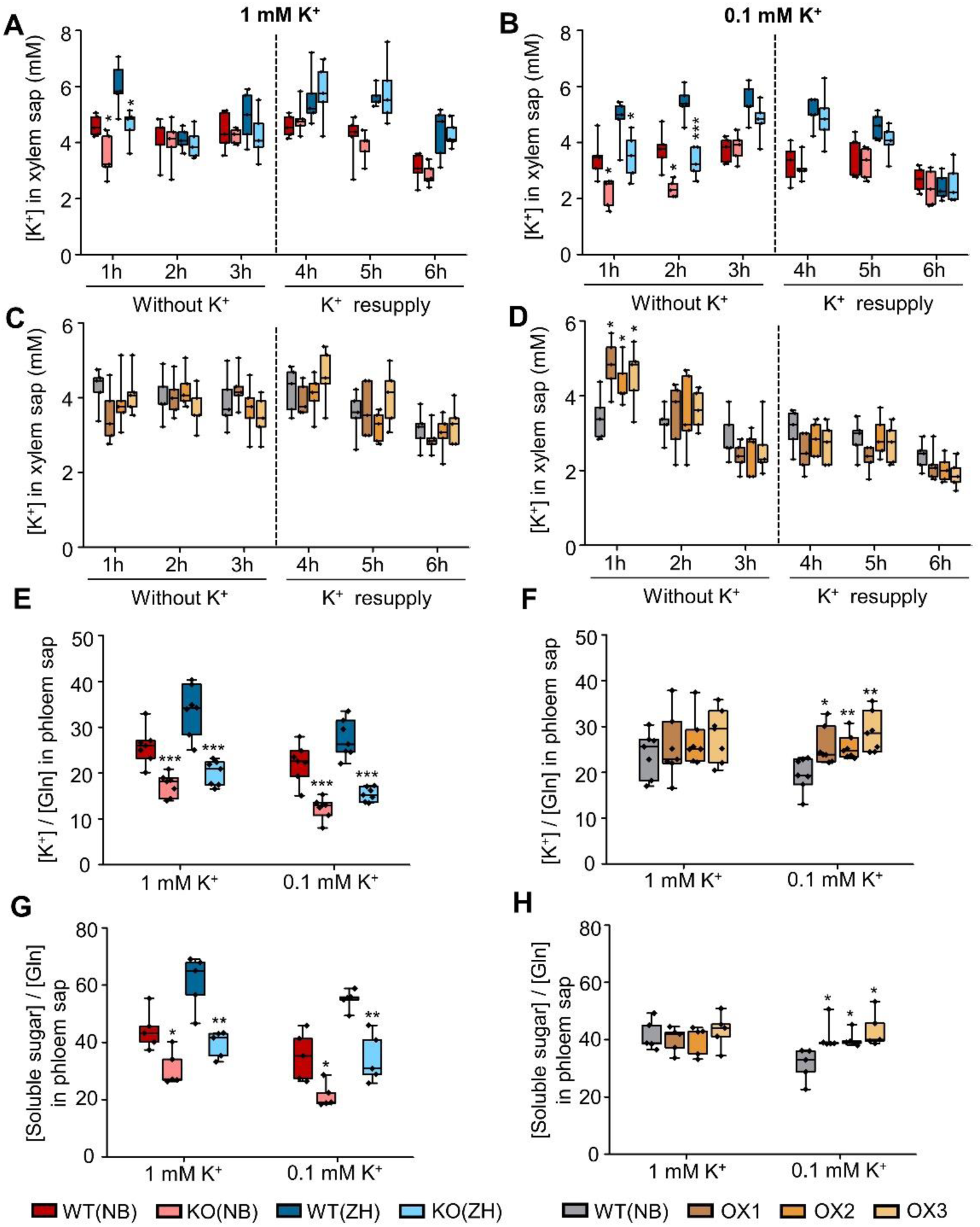
Effect of *OsHAK18* expression level on K^+^ concentration in the xylem and soluble sugar concentration in phloem sap of rice. A-D, The seedlings were grown in IRRI solution containing 1 mM K^+^ for 8 weeks, then transferred to a medium with 1 mM K^+^ (A and C) or 0.1 mM K^+^ (B and D) for 3 days. Subsequently, the shoot was excised, the root was transferred to 0.1 mM CaCl_2_ solution without K^+^ and the xylem sap was collected during three one-hour periods. After 3 hours, the root was transferred back to 1 mM K^+^ (A and C) or 0.1 mM K (B and D) solution and the xylem sap was collected similarly, in three one-hour periods for 3 more hours. E-H, The seedlings were grown in IRRI solution containing 1 mM K^+^ for 8 weeks, then transferred to a medium with 1 or 0.1 mM K^+^. After 3 days of culture, phloem sap was collected during a 5 h period for K^+^ and soluble sugar analysis. Glutamine was determined for internal standard (Ren *et al*., 2005). The box plots and t-tests are as in Figure 2. Significant differences from the corresponding WT are indicated by asterisks (*, P<0.05; **, P<0.01; ***, P<0.001).

In the OX lines grown in 1 mM K^+^, the xylem sap K^+^ concentration was the same as in WT and did not change with time (Figure 4C), matching the pre-experiment lack of difference between WT’s and OX‘s root [K^+^] (Figure 3F). In contrast, in OX grown in 0.1 mM K, root [K^+^] was higher than in WT during the first h of K^+^ deprivation and declined to the same level as WT’s during the next two hours (Figure 4D). Thus, also the root [K^+^] of the OX plants required an equilibration period. After K^+^ was added back, K^+^ concentration in the xylem sap of KO or OX lines remained similar to their corresponding WTs and did not increase during the three hs of the follow-up period (Figure 4, A-D), again, excluding OsHAK18’s role in xylem sap loading with K^+^.

### OsHAK18 is involved in K^+^ re-circulation and affects sugar translocation in the phloem

Based on the above results, it is reasonable to assume that OsHAK18 influences K^+^ root and shoot distribution via phloem transport. To examine this directly, phloem sap was collected from all our rice genotypes grown in a medium with 1 mM or 0.1 mM K^+^. Because glutamine is abundant in the phloem sap and remains quite constant during the entire 24 hs day-night cycle, glutamine is often used as an internal standard for measuring the concentration of ion and photo-assimilates in phloem sap (Corbesier et al., 2001; Ren et al., 2005). The phloem sap [K^+^] was thus normalized to the glutamine concentration in the same sample. The phloem sap [K^+^] in both KO lines was only about 50%-65% of that in the corresponding WTs irrespective of the [K^+^]_EXT_ levels (Figure 4E), while in all three OX lines in grown in [K^+^]_EXT_ of 0.1 mM it increased by at least 30% (Figure 4F). These results support the notion that OsHAK18 contributes to K^+^ entry into the phloem.

Since translocation of photo-assimilates from source leaves to roots via phloem requires high K^+^ concentrations in the sieve tubes and a steep transmembrane pH gradient, we examined the OsHAK18 effect on sugar concentration in the phloem. Remarkably, KO and OX of *OsHAK18* affected oppositely the soluble sugar concentration in the phloem at the low [K^+^]_EXT_: KO reduced and OX increased it (Figure 4, G and H). In contrast, at the normal K^+^ supply level, while KO reduced the sugar level, OX did not affect it (Figure 4, G and H). These changes in the phloem sugar level were consistently similar to changes in the phloem [K^+^] of these transformants.

To test further whether OsHAK18 contributes to K^+^ recycling from shoot to root, we took advantage of the Rb^+^ transport capability of the HAK family transporters (Ródenas et al., 2017) and performed an experiment with split-root plants. One half-root was first exposed to 0.5 mM Rb^+^ and the other half – to 0.5 mM K^+^ for 3 hs. After that, the medium in both root containers was switched to one with 0.5 mM K^+^ (Figure 5A). We measured the Rb^+^ content in both parts of the root and the shoot at 3 hs and 6 hs after initiation of the treatments.

**Figure 5.**
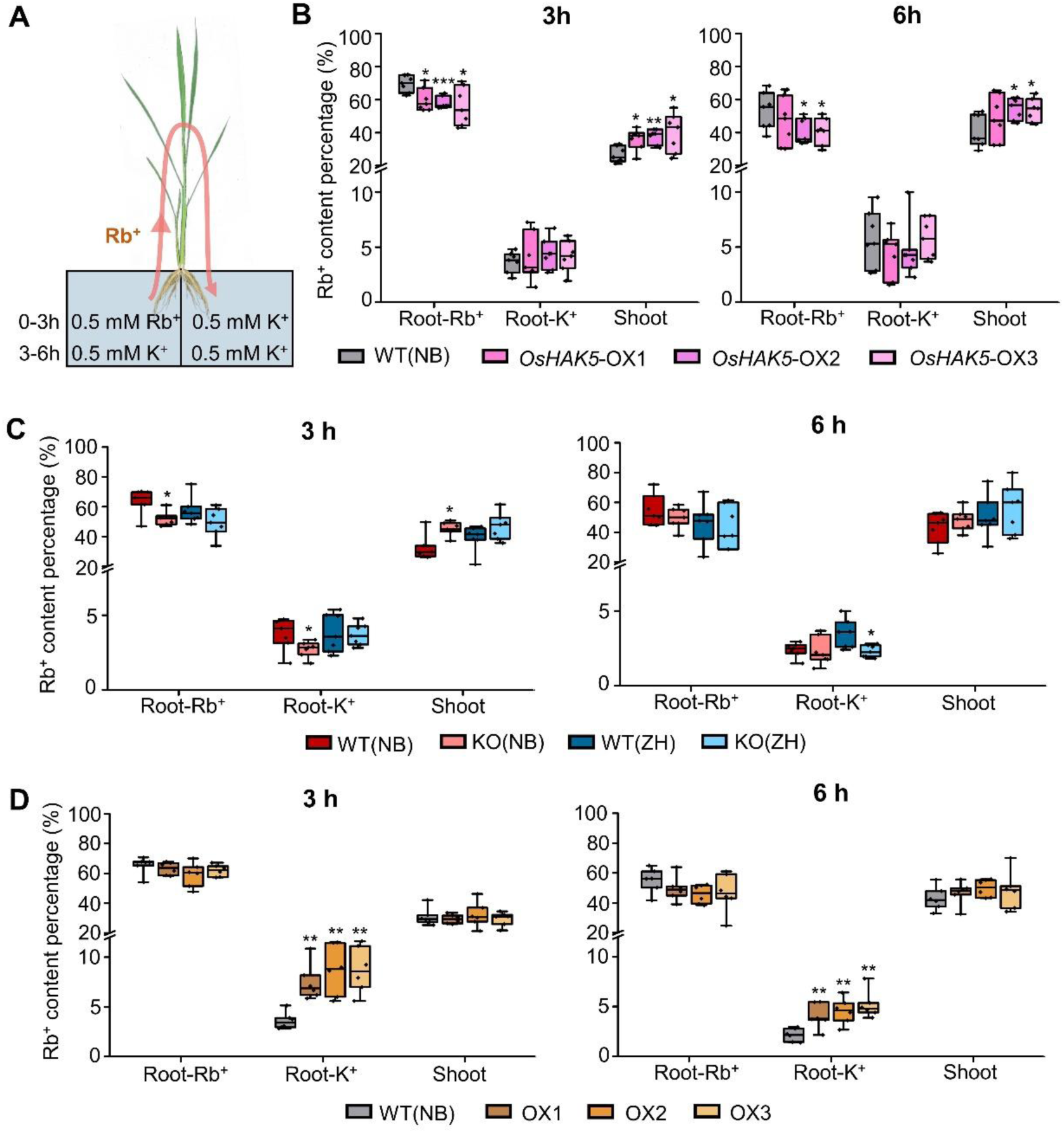
Effect of *OsHAK18* expression level on Rb^+^ transport in rice plants. A, Schematic diagram of split-root experiment. B, Rb^+^ content percentage (i.e., the percentage of the total Rb^+^ acquired by the plant) in different plant parts in *OsHAK5* overexpression lines and their corresponding WT at 3 h and 6 h of the pulse chase. C, *OsHAK18* KO lines and their corresponding WTs. D, *OsHAK18* overexpression lines and their corresponding WT. Root-Rb^+^, Root exposed to Rb^+^; Root-K^+^, Root not exposed to Rb^+^. The box plots and t-tests are as in Figure 2. Significant differences from the corresponding WT are indicated by asterisks (*, P<0.05; **, P<0.01; ***, P<0.001).

As a control, we performed the same experiment on rice overexpressing *OsHAK5*, a transporter known to mediate root to shoot K^+^ translocation in rice. In *OsHAK5*-OX, as expected, the relative amount of Rb^+^ in the Rb^+^-exposed half root (i.e., the percentage of the total Rb^+^ acquired by the plant) was lower than in WT, while in the roots exposed to K^+^, the relative amount of Rb^+^ was the same as in WT (Figure 5B). In the shoot, Rb^+^ accumulation was enhanced in *OsHAK5*-OX relative to WT (Figure 5B). The fulfilled expectation with regard to the *OsHAK5*-OX effect, together with an earlier demonstration of Rb^+^ transport by OsHAK18 (Figure 2C), confirmed that the Rb^+^ distribution in the split-root experiment with OsHAK18 should also reliably represent the circulation of K^+^ in rice.

The relative amount of Rb^+^ in the Rb^+^-exposed half root was almost the same in both *OsHAK18*-KO lines and their corresponding WTs (roughly 50-70 %) except that in the NB KO line it was less than in NB WT in 3h of treatment. The root of the K^+^-exposed half contained a very low fraction of Rb^+^ (roughly 4% of total), KO reduced this even further (by roughly 25-30% of WT, the NB cultivar after 3 hs and the ZH cultivar after 6 hs; Figure 5C). In the shoot, which accumulated about 40% of the Rb^+^ in 3 hs and 50% Rb^+^ in 6 hs, the NB KO line accumulated more Rb^+^ than NB WT in 3h of treatment (Figure 5C). In contrast, the Rb^+^ content in the K^+^-exposed roots of *OsHAK18*-OX plants was more than double that in WT at both hour 3 and hour 6 (Figure 5D). All these data support the notion that the direction of K^+^ flow mediated by OsHAK18 is opposite from that of OsHAK5, and that OsHAK18 enhances K^+^ translocation from shoot to root.

### Increasing *OsHAK18* expression affects the plant architecture and the sugar translocation resulting in an increased yield of rice in a paddy field

To reveal the role of OsHAK18 in the rice maturation stage, we evaluated the plant growth and the grain yield of WT, KO, and OX mature (20-weeks old) plants. Compared to WT, *OsHAK18*-KO plants had appreciably reduced height, tiller number, panicle length, seed setting rate (%) and grain number per panicle, leading to reduced yield and grain-to-straw ratio (Figure 6, A and B; Supplemental Figure S8, A and C). In contrast, while *OsHAK18*-OX plants had also reduced plant height, they had markedly higher tiller numbers and therefore, a higher grain yield (Figure 6, F and G; Supplemental Figure S8, B and D).

**Figure 6.**
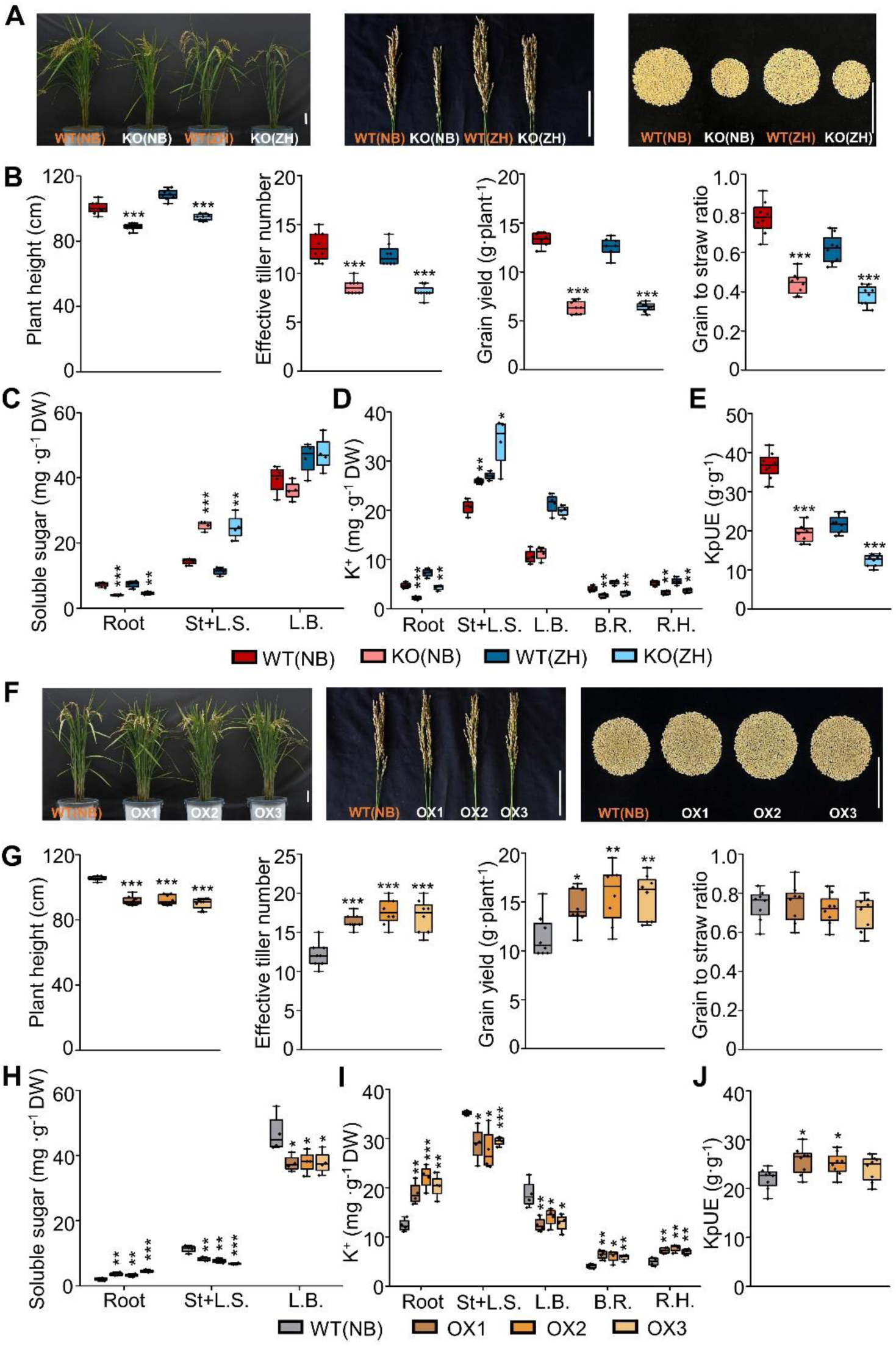
Effect of *OsHAK18* expression level on agronomic traits and K^+^ and soluble sugar distribution at reproduction. Rice plants were grown in the paddy field and irrigated, fertilized and medicated regularly until they were fully mature (approx. 20 weeks old). A and F, Phenotypes of whole plants and panicles, and yield per plant. Scale bars = 10 cm. B and G, Plant height, effective tiller number, grain yield, grain to straw ratio. C and H, Soluble sugar concentration. D and I, K^+^ concentration. E and J, KpUE, K^+^ physiological utilization efficiency, i.e., the grain yield produced by unit weight of total accumulated K^+^ per plant. St+L.S., Stem and leaf sheath; L.B., Leaf blade; B.R., Brown rice; R.H., Rice hull. Detailed data on other agronomic traits are provided in Supplemental Figure S8. The box plots and t-tests are as in Figure 2. Significant differences from the corresponding WT are indicated by asterisks (*, P<0.05; **, P<0.01; ***, P<0.001).

Since K^+^-generated turgor pressure in the phloem is essential for the loading of soluble sugar at the source and unloading at sink sites (Dryer et al., 2017), we asked whether OsHAK18 had a role in these processes. Therefore, we examined the soluble sugar distribution in the root, stem, leaf sheath and leaf blade. In the *OsHAK18*-KO plants, soluble sugar accumulation in the root was markedly lower compared to WT, but in the stem and leaf sheath, it was higher than in the WT (Figure 6C). In contrast, in the *OsHAK18*-OX plants soluble sugar content was higher in the roots but lower in the stem, leaf sheath, and leaf blade in comparison with the WT (Figure 6H).

To further outline the OsHAK18 role in altering K^+^ distribution between source and sink organs at the maturation stage, we measured K^+^ concentration in the root, stem, leaf sheath, leaf blade, brown rice (a whole rice grain without the hull), and rice hull. As expected, K^+^ concentration in the root, brown rice and rice hull was reduced by *OsHAK18*-KO and increased by *OsHAK18*-OX. In contrast, K^+^ concentration in stem and leaves was increased by *OsHAK18*-KO and decreased by *OsHAK18*-OX (Figure 6, D and I). The differences in K^+^ concentration among the samples were consistent with those of the sugar. Interestingly, K^+^ physiological utilization efficiency (KpUE), i.e., the grain yield normalized (w/w) to the total accumulated K^+^ in a plant, was almost halved by *OsHAK18*-KO (Figure 6E), but increased by 9% - 17% by *OsHAK18*-OX (Figure 6J), demonstrating that OsHAK18-controlled K^+^ re-circulation in the phloem is essential for maintaining a high KpUE. Thus, properly enhancing the OsHAK18 activity driven by its endogenous promoter can promote both grain yield and K^+^ use efficiency.

### OsHAK18 mediates Na^+^ recycling from shoot to root under salt stress

Since OsHAK18 enhanced also the accumulation of Na**^+^** in yeast (Figure 2C), we examined its function in rice under 100 mM NaCl stress. Relative to WT, *OsHAK18*-KO reduced both K^+^ and Na^+^ concentration in the root, while increasing the concentration of Na^+^, but not K^+^, in the shoot (Supplemental Figure S9, A and C). Their calculated K^+^/Na^+^ ratio was lower than in WT in both root and shoot (Supplemental Figure S9E). In contrast, *OsHAK18*-OX elevated K^+^ concentration but decreased Na^+^ concentration in the shoots with the opposite change of K^+^ and Na^+^ concentration in the root (Supplemental Figure S9, B and D), and thus resulted in a higher K^+^/Na^+^ ratio compared to WT in the shoot (Supplemental Figure S9F).

Ion recycling under high salt stress may be affected due to a disruption of phloem transport by plasmodesmata plugged by callose accumulation and P-protein aggregation at the sieve pores (Xie and Hong, 2011). To distinguish this effect of the salt stress on plasmodesmata transport from the effect of salt stress on membrane transport via OsHAK18, we treated all our rice genotypes with a moderate salt stress of 20 mM Na^+^ (at [K^+^]_EXT_ of 1 mM) for 3 days and measured K^+^ and Na^+^ in the phloem sap. Both K^+^ and Na^+^ concentrations were appreciably lower in the sap of the *OsHAK18*-KO plants than in WT (Figure 7, A and B), and markedly higher (by about 30-40 %) in the phloem sap of the *OsHAK18*-OX lines compared to WT (Figure 7, C and D). The results suggest that under salt stress, in rice, OsHAK18 enhances Na^+^ (K^+^) recycling from the shoot to the root.

**Figure 7.**
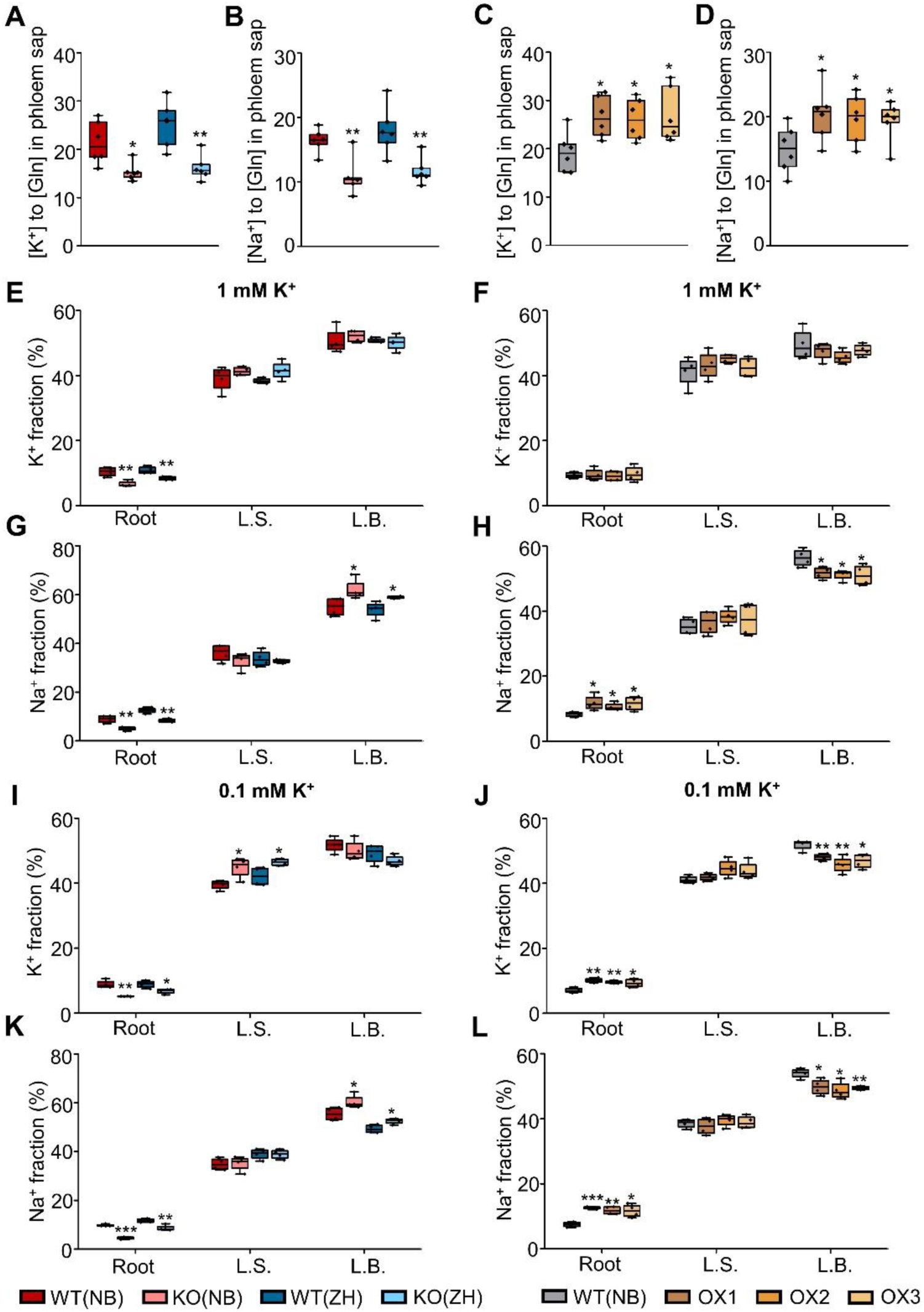
Effect of *OsHAK18* expression level on K^+^ and Na^+^ transport and distribution in rice supplied with NaCl in the culture solution. A-D, Ten-day-old seedlings were grown in IRRI solution containing 1 mM K^+^ for 8 weeks, then transferred to a solution containing additionally 20 mM NaCl and allowed to grow for 3 d. The phloem sap was collected and analyzed. E-L, Ten-day-old seedlings were grown in IRRI solution containing 1 mM K^+^ for 2 weeks, then transferred to 1 mM K^+^ (E-H) or to 0.1 mM K^+^ (I-L) solutions for 2 weeks, then transferred to 1 mM K^+^ or to 0.1 mM K^+^ solutions containing additionally 20 mM NaCl and allowed to grow for 3 d, then returned to 1 mM or 0.1 mM K^+^ medium without NaCl for additional 3 d. K^+^ or Na^+^ fraction (%) represents the percentage of K^+^ or Na^+^ content in each plant part of the total K^+^ or Na^+^ content of the whole plant. L.S., Leaf sheath; L.B., Leaf blade. Detailed data on K^+^ and Na^+^ concentration in root and shoot are provided in Supplemental Figure S10. The box plots and t-tests are as in Figure 2. Significant differences from the corresponding WT are indicated by asterisks (*, P<0.05; **, P<0.01; ***, P<0.001).

In a follow-up experiment, we treated all rice lines with 20 mM NaCl for 3 days at [K^+^]_EXT_ of 1 mM and 0.1 mM. We returned the plants to 1 mM and 0.1 mM [K^+^]_EXT_ medium without NaCl for additional 3 days and then measured K^+^ and Na^+^ content in the roots and shoots. At [K^+^]_EXT_ of 1 mM, although K^+^ concentration and its relative distribution in shoots of KO and OX plants were similar to those of WT (Supplemental Figure S10, A and B; Figure 7, E and F), Na^+^ concentration in the leaf sheath and the leaf blade of KO plants was markedly higher than in the respective WT plants (Supplemental Figure S10C). Furthermore, compared to WT, *OsHAK18*-KO plants had a lower fraction (% of total accumulated amount per plant) of Na^+^ in the roots but a higher Na^+^ fraction in the leaf blade (Figure 7G), and these differences were exactly opposite in the *OsHAK18*-OX plants (Figure 7H). These differences resulted in a decrease of K^+^/Na^+^ ratio in the shoot of *OsHAK18*-KO and an increase in the shoot of *OsHAK18*-OX.

At [K^+^]_EXT_ of 0.1 mM, the concentration of K^+^ and Na^+^ in the roots of *OsHAK18*-KO plants was lower than in the corresponding WT plants, while K^+^ concentration in the leaf sheath and Na^+^ concentration in both the leaf sheath and the leaf blade of KO plants were markedly higher than in the corresponding WT plants (Supplemental Figure S10, E and G). Therefore, compared to WT, *OsHAK18*-KO plants had a lower fraction of K^+^ and Na^+^ in the roots but a higher K^+^ fraction in the leaf sheath as well as a higher Na^+^ fraction in the leaf blade (Figure 7, I and K). Moreover, these differences were exactly the opposite in the *OsHAK18*-OX plants with regard to Na^+^ (Supplemental Figure S10 H compared to G, Figure 7 L compared to K), and also with regard to K^+^ (with a “switch” between L.S. vs. L.B.; Supplemental Figure S10 F compared to E, Figure 7 J compared to I). All these results indicate that under salt stress, OsHAK18 contributes to the recirculation of Na^+^ as well as of K^+^ from shoot to root, and this is more pronounced at the limited than at the regular K^+^ supply.

## DISCUSSION

That K^+^ circulates in plants from root to shoot and then back from shoot to root has been already textbook knowledge for years. Out of the rice HAK/KUP/KT genes of K^+^ carriers, the *OsHAK18* gene product, OsHAK18, is the first of its family to be shown here unequivocally to mediate the downward, shoot-to-root K^+^ translocation via the phloem, and translocation of Na^+^ under the presence of Na^+^ salts. Enhancing *OsHAK18* expression controlled by its native promoter can dramatically promote sugar translocation, grain yield and K^+^ physiological utilization efficiency in rice (Figure 6, F-J).

The specific cellular localization of OsHAK18 suggests its role in several stages of this K^+^ recirculation (the following description of its activity is based solely on OsHAK18’s expression patterns, Figure 1): In source leaves, OsHAK18 mediates the transport of K^+^ from MCs to the sieve element-companion cell (SE-CC) complex (Figure 8A). Generally, the short-distance transport of K^+^ in source leaves may be achieved through two distinct pathways (Figure 8A, dashed horizontal arrows) and/ or their combination: (i) K^+^ leaked to the apoplastic space from mesophyll cells (MCs) can be imported by OsHAK18 into the adjacent MCs or into the adjacent vasculature bundle sheath (VBS) cells, then diffuse via plasmodesmata into mestome sheath cells (MS) cells, then, it can leak again into the apoplast (including the xylem) and be imported again by OsHAK18 (stained in Figure 1Dc and especially strongly in Figure 1Cc) into the adjacent xylem parenchyma cells (XPCs), then it can leak again into the apoplast and be imported by OsHAK18 into phloem parenchyma cells/ companion cells (PPCs/CCs), and then diffuse into phloem sieve elements through plasmodesmata; (ii) at the other extreme, not involving membrane transporters, K^+^ can diffuse all the way from mesophyll cells (MCs) to the phloem sieve elements via plasmodesmata (symplastically), through the vasculature bundle sheath (VBS) cells and mestome sheath (MS) cells, and various parenchyma cells (incl. phloem parenchyma cells/ companion cells (PPCs/CCs). Notably, as interfering with OsHAK18 expression shows (see discussion further down), the purely symplastic pathway is probably much less effective for the downward translocation of K^+^.

**Figure 8.**
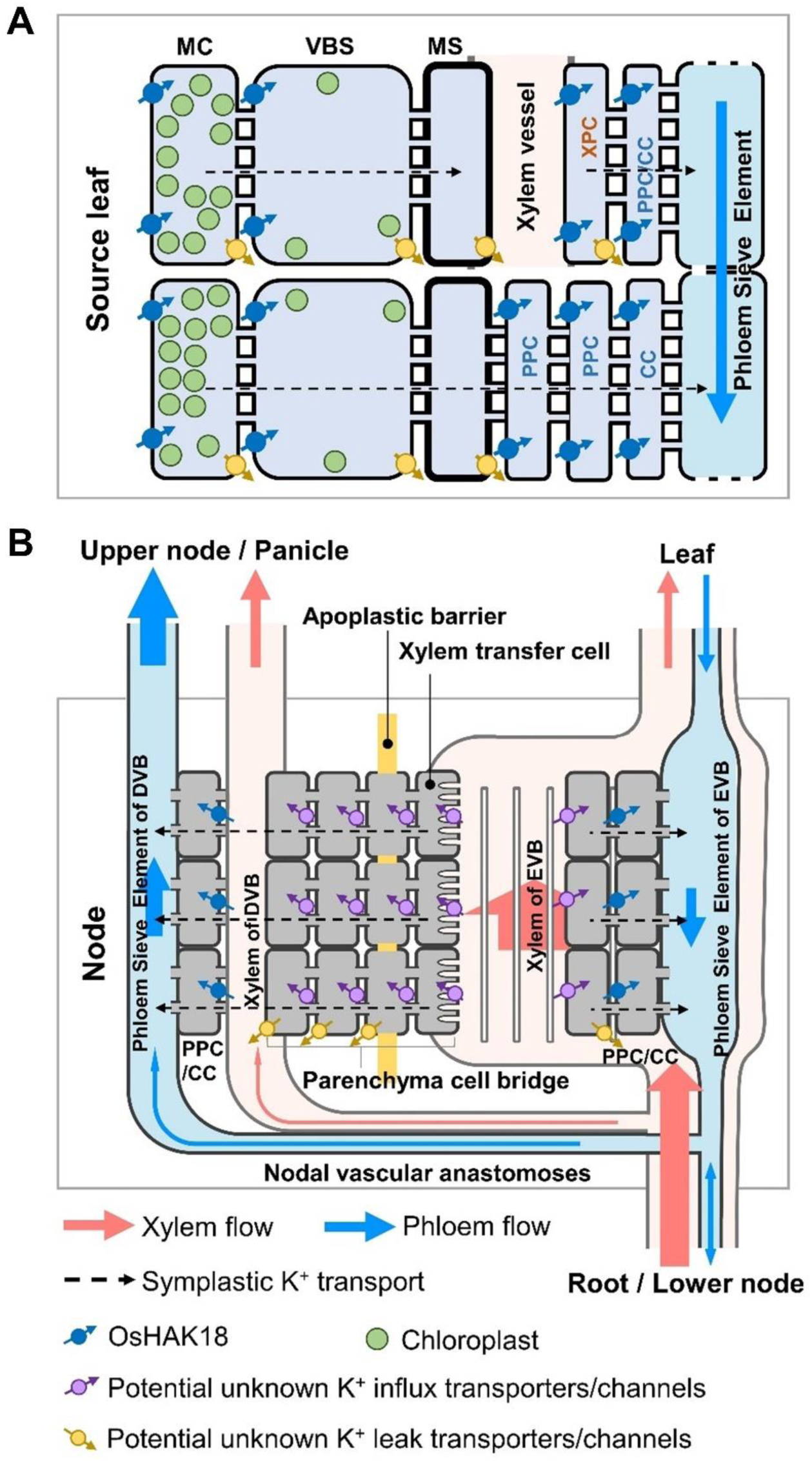
A working model for the roles of OsHAK18 in K^+^ redistribution in the leaf blades and nodes of rice. A, The role of OsHAK18 in the short-distance transport of K^+^ in a source leaf. B, The role of OsHAK18 in a node. CC, companion cell; DVB, diffuse vascular bundle; EVB, enlarged vascular bundle; MC, mesophyll cell; MS, mestome sheath cell; PPC, phloem parenchyma cell; VBS, vascular bundle sheath cell; XPC, xylem parenchyma cell. The size of arrows in the phloem and xylem represents flow area.

It should be noted that phloem bulk flow in rice is not only downward, but also conveys minerals and photo-assimilates upwardly, from the source leaves to the sink leaves and organs; the branching into the two types of flow occurs in the node. Thus, in the rice node vascular system (Figure 8B, referring to the schematic diagram by Yamaji and Ma, 2017), OsHAK18 mediates the influx of K^+^ into PPCs/CCs of the enlarged vascular bundle (EVB) and of the diffuse vascular bundle (DVB; Figure 8B). In future work, it would be of interest and significance to identify the different K^+^ efflux and K^+^ influx transporters in both leaves and nodes.

The evidence for OsHAK18 function in K^+^ recirculation via the phloem is based on an ensemble of nine findings using a variety of approaches (all the comparisons are in relation to WT): (1) the natural *OsHAK18*-enriched expression in the phloem CCs and PPCs and particularly in the shoot (Figure 1, B-D); (2) the membrane localization of OsHAK18 in rice protoplasts (Figure 2, A and B); (3) a decreased [K^+^] in the root of *OsHAK18-KO* in both Nipponbare and Zhonghua11 cultivars (Figure 3, A and C); (4) the decreased [K^+^] in the leaf blade of *OsHAK18*-OX in the Nipponbare cultivar plants exposed to K^+^-deficient solution (Figure 3D); (5) the increased shoot-to-root ratio of total K^+^ content in the KO plants (Figure 3, E and G), and the decreased shoot-to-root ratio of total K^+^ content in *OsHAK18*-OX plants faced with K^+^-deficient nutrient solution (Figure 3H); (6) identical rates – among all genotypes within a given group (KO or OX) – of K^+^ depletion from the media bathing the roots, independent of both K^+^ supply levels (low and high), even if 3-fold slower at the low [K+]_EXT_). These findings exclude OsHAK18 as a root K^+^ acquisition transporter (Supplemental Figure S7); (7) a reduced [K^+^] in the phloem sap of *OsHAK18*-KO plants (Figure 4E; Figure 7, A and B) and increased [K^+^] in the phloem sap of *OsHAK18*-OX plants (Figure 4F; Figure 7, C and D); (8) the decreased Rb^+^ translocation from the Rb^+^-fed root half to the Rb^+^-spared root half in the split root experiment in the KO plant (Figure 5C) and the enhanced Rb^+^ translocation from the Rb^+^-fed root part to the Rb^+^-spared root part in the OX lines (Figure 5D); (9) the diametrically opposed to the above distribution of K^+^ between the root parts of *OsHAK5*-OX in the control split-root experiment (Figure 5B), due to the known function of OsHAK5 in enhancing translocation of K^+^ from root to shoot (Yang et al., 2014; 2020).

### OsHAK18 is not involved in root K^+^ absorption nor in root-to-shoot K^+^ translocation

Root absorption of K^+^ from culture medium is mediated by high- and low-affinity transporters and channels which enable adaptation to varied supply levels (Epstein et al., 1963; Maathuis and Sanders, 1997; Li et al., 2017). Root-acquired K^+^ is transported into root stelar tissues and translocated from the root toward the shoot via xylem vessels (Marschner et al., 1996). As detected in Arabidopsis and rice, most of the putative HAK transporters and K^+^ channels are located in root epidermal and xylem parenchyma cells, indicating their function mainly in root K^+^ absorption and K^+^ translocation to the shoot. In Arabidopsis, AtHAK5 is the only system for root K^+^ uptake from as low as 0.01 mM external [K^+^] (Rubio et al., 2010), and AtKUP7 shows a higher expression level in root than in other organs and is crucial for K^+^ uptake (Han et al., 2016). In rice, both OsHAK1 and OsHAK5, which have the highest homology with AtHAK5, participate in K^+^ absorption and are up-regulated by K^+^ deficiency in roots (Yang et al., 2014; Chen et al., 2015). *OsHAK16* was also up-regulated by K^+^ deficiency in roots and plays a similar but non-redundant role in K^+^ uptake by roots as do OsHAK1 and OsHAK5 (Feng et al., 2019). In this study, we found that, unlike those already-described HAK-family members, *OsHAK18* is expressed mainly in the phloem CCs, but not in epidermal and cortex cells of the root (Figure 1A), and its transcripts in the root are not up-regulated by K^+^ deficiency (Supplemental Figure S2A).

While in the long-term experiments, *OsHAK18*-KO decreased the [K^+^] in the root and increased [K^+^] in the shoot (Figure 3, A and C), the lack of effect of *OsHAK18* elimination on the short-term K^+^ absorption by the root (Supplemental Figure S7, A and B), indicates that OsHAK18 is not involved in root K^+^ uptake in rice. This conclusion is further strengthened by the *OsHAK18*-OX under its native promoter that we generated to enhance OsHAK18 activity without affecting its pattern of expression and localization to avoid potential alteration of its K^+^ transport properties. As expected, *OsHAK18*-OX did not affect the short-term K^+^ absorption rate (Supplemental Figure S7, C and D). A similar conclusion is also suggested by the lack of effect of either *OsHAK18*-KO or *OsHAK18*-OX on the K^+^ concentration in the xylem sap (compared to the corresponding WT), irrespective of the external K^+^ supply levels at the root (Figure 4, A-D).

### OsHAK18 promotes K^+^ circulation from shoot to root and sugar translocation from source to sink organs

In Arabidopsis, a “weakly-inward rectifying” K^+^ channel, AKT2, has been shown to mediate K^+^ phloem loading, but also (under different circumstances) to mediate K^+^ release from the phloem, thereby enhancing the loading of phloem with sugar (Deeken et al., 2002, Gajdanowicz et al., 2010). Similar to AKT2, OsAKT2 has been shown to be important for K^+^ phloem loading and K^+^ recirculation via phloem (Tian et al., 2021). However, it was not known if additional K^+^ transporter(s) is also involved in K^+^ movement in the phloem, especially in the K^+^ translocation from shoot to root. The fact that *OsHAK18*-KO plants are impaired in K^+^ recirculation means that OsAKT2 cannot fulfill this role all by itself. OsHAK18 has now been revealed as a new participant in the phloem loading of K^+^ and in the shoot-to-root translocation of K^+^. Furthermore, OsHAK18’s participation in sugar translocation paralleling with K^+^ redistribution (Figure 4, G and H; Figure 6, C and H) resembles the function of OsAKT2 which enhances sugar transfer from source sites to sink sites (Tian et al., 2021), demonstrating that OsHAK18 coordinates the delivery of K^+^ and photo-assimilates in the phloem.

OsAKT2 and OsHAK18 may play complementary roles in the phloem movement of K^+^ at different K^+^ supply levels or different loading directions. On one hand, OsAKT2 functions mainly as an inward rectifier with strong voltage dependency (Huang et al., 2021) and the elimination of phloem-located *OsAKT2* expression resulted in relatively higher accumulation of K^+^ in the shoot at a normal K^+^ supply (Tian et al., 2021). Notably, OsHAK18 contributes to K^+^ re-circulation in phloem at both normal and low K^+^ supply status (Figure 3, E and H; Figure 4, E and F). In addition, when expressed in Arabidopsis, OsAKT2 can complement the K^+^ deficiency in the phloem sap and the leaves of *akt2* mutant (Huang et al., 2021). On the other hand, however, unlike AKT2 which functions in the leakage of K^+^ from the companion cells of the transport phloem (Deeken et al., 2002; Gajdanowicz et al., 2010), OsAKT2 does not allow K^+^ leak from cells (Huang et al., 2021). Furthermore, since this outward K^+^ leak is what allows the Arabidopsis AKT2 to generate the sugar-loading-into-phloem “K^+^ battery” (Gajdanowicz et al., 2010), the lack of outward K^+^ leak via OsAKT2 casts doubt on its participation in any such battery in rice (Huang et al., 2021). Therefore, it seems that in the rice plant, OsAKT2 and OsHAK18 share together the responsibility for K^+^ loading into the phloem, but between the two, only OsHAK18 is likely to be involved in the phloem sugar loading, perhaps “partnering” instead with a dedicated K^+^ release channel, such as OsGORK (that is expressed in rice more widely than its homolog, OsSKOR; Kim et al., 2015).

### OsHAK18 activity manifests differently at different K^+^-supply levels

The OsHAK18 relevance for K^+^ recirculation, which was not evident in the WT, becomes obvious in *OsHAK18*-KO, (a) as a decrease of [K^+^] in the root (relative to WT), irrespective of K^+^ supply ([K^+^]_EXT_; Figure 3, A and C), and (b) as an increase of [K^+^] in the shoot (relative to WT), but only at the low [K^+^]_EXT_ (Figure 3C).

The decrease in the root’s [K^+^] in the KO is consistent with a decline in K^+^ return from the shoot, due to a decreased K^+^ phloem loading in the absence of *OsHAK18*, attesting to its non-negligible role in the WT. In support of this notion, at [K^+^]_EXT_ of 1 mM, the phloem sap [K^+^] was indeed appreciably lower in the *OsHAK18*-KO than in the WT (Figure 4E). This difference held also at the low [K^+^]_EXT_ (at least, after three days of growth at [K^+^]_EXT_ of 0.1 mM K^+^; Figure 4E).

Furthermore, given that *OsHAK18*-KO did not affect the rates of K^+^ upward movement in the xylem, irrespective of [K^+^]_EXT_ (Figure 4, A and B, hours 4-6; Supplemental Figure S7, A and B), in the absence of this downward-translocating transporter, the shoot’s [K^+^] would be expected to increase. Why, then, did the shoot’s [K^+^] increase in KO only at [K^+^]_EXT_ of 0.1 mM (Figure 3C)? This too can be explained by the decreased K^+^ return via the KO’s phloem. That it is not reflected in the shoot’s [K^+^] at [K^+^]_EXT_ of 1 mM, is probably due to the large K^+^ storage capacity of the shoot, particularly in the leaf sheath, which at the normal K^+^ supply, buffers small changes in [K^+^].

Moreover, why is the root’s [K^+^] decline in the KO plants at [K^+^]_EXT_ of 1 mM and 0.1 mM (due to diminished downward K^+^ return), mirrored in the OX plants as the root’s K^+^-enrichment only at [K^+^]_EXT_ of 0.1 mM (Figure 3D),but not at [K^+^]_EXT_ of 1 mM (Figure 3B)? The same question applies to the OX shoot’s [K^+^], which shows depletion (in the leaf blade) only at [K^+^]_EXT_ of 0.1 mM (Figure 3D). It would appear that the increased expression of *OsHAK18* (under its native promoter; Supplemental Figure S4F) is not effectively pronounced at the transport activity level at the normal K^+^ supply, perhaps because of its post-translational modification. In support of this notion of the lack of added OsHAK18 activity in OX at the normal K^+^ supply, is also the unaltered [K^+^] in the phloem sap (Figure 4F).

However, at [K^+^]_EXT_ of 0.1 mM, *OsHAK18* expression in the shoot may be even enhanced (resembling its enhanced expression on day 3 of root treatment with 0 mM K^+^; Supplemental Figure S2, B and C). Since such expression enhancement, and possibly also post-translational modification, would likely include the additional gene copies under the *OsHAK18*’s native promotor in OX, OsHAK18 would play a prominent role in re-circulating K^+^ from the shoot to the root (Figure 3D; Figure 4F). Moreover, the fact that lowering the K^+^ supply level emphasized the effect of OsHAK18 OX on [Na^+^] in the root, but even more so in the shoot (Supplemental Figure S10 H compared to D), supports the notion of an increased activity of OsHAK18 also toward Na^+^, as a result of a regulatory feedback of the K^+^ level on the transporter itself. By comparison, also OsHAK5’s role as an upward K^+^-translocating transporter, is similarly not pronounced at [K^+^]_EXT_ of 1 mM, but becomes significant at [K^+^]_EXT_ of 0.1 mM (Yang et al., 2014).

### Enhancing OsHAK18 expression increases K^+^ physiological utilization efficiency and grain yield

Improving K^+^ physiological use efficiency, i.e., less K^+^ demand for reaching high yield, can reduce soil K^+^ depletion by plants and the demand for K^+^ fertilizers (Shin, 2014). Unlike N and P which are mainly accumulated in grains/seeds, the majority of acquired K^+^ from soil or fertilizer is kept in vegetative organs, like straw, in rice. The dramatic reduction of K^+^ concentration in the straw of *OsHAK18*-OX (Figure 6I) and its increased grain yield (Figure 6G) signify an increase of K^+^ physiological utilization efficiency by a non-negligible 9-17% (Figure 6J), resulting – we presume – from an enhanced circulation of K^+^ in the phloem.

We noticed that OsHAK18 had a significant effect on rice architecture, especially tiller number which is one of three major components in determining the rice grain yield (Figure 6, B and G; Supplemental Figure S8). Plant architecture including rice tiller number is regulated by polar auxin transport (Lu et al., 2015; Luo et al., 2020). Cellular auxin fluxes are strongly affected by transmembrane pH gradients (Steinacher et al., 2012), which, in turn, are regulated by interactions between H^+^ pumps, H^+^-coupled carriers and ion channels at the PM (Sze and Chanroj, 2018). For example, OsHAK5, an electrogenic K^+^/H^+^ symporter, regulated the chemiosmotic gradient, PM polarization, and extracellular pH in rice, resulting in positive regulation of polar auxin delivery streams and an increased rice tiller number (Yang et al., 2020). Since OsHAK18 is a plasma-membrane-located K^+^/H^+^ co-transporter (Figure 2), expressed abundantly in the vascular tissue (Figure 1B), we speculate that OsHAK18 affects the pH in the apoplast, and thus, among others, controls tiller emergence in the basal node, thus affecting rice architecture (Figure 6, B and G). These findings may in the future render OsHAK18 a target for manipulations aimed at increasing the rice yield and K^+^ use efficiency in the field.

## MATERIALS AND METHODS

### Plant materials and growth conditions

Two cultivars of rice (*Oryza sativa* subsp. *japonica*) were used in this work, Nipponbare (NB) and Zhonghua11 (ZH). For hydroponic experiments, seed sterilization and basal nutrient solution composition for seedling growth were described previously (Ai et al., 2009). Similar-size ten-day-old rice seedlings of each wild-type (WT) and transgenic lines were selected and transferred to IRRI nutrient solution (Yoshida et al., 1976; Yang et al., 2014). For K^+^ starvation or depletion experiments, rice seedlings were grown in the IRRI solution containing 1 mM K^+^ for 4 weeks and then transferred into a solution in which KH_2_PO_4_ and K_2_SO_4_ were replaced by NaH_2_PO_4_ and KCl, respectively. For salt stress experiments, 100 mM NaCl was added in the IRRI solution after 4 weeks. In all treatments, the solutions were replaced every 2 days. The hydroponic experiments were carried out in a greenhouse with a 16h-light/8h-dark, temperature between 22-30°C, and 60% to 70% relative humidity. For field experiments, rice plants were grown in paddy soil with regular fertilization during the normal growing seasons in Nanjing, China. At each harvest, roots of all rice plants were washed in 0.1 mM CaSO_4_ for 5 min. Roots, stems, leaf sheaths and blades, brown rice and rice hull were separated before their biomass was recorded and concentrations of K^+^ and Na^+^ were measured as described previously (Ding et al., 2006; Yang et al., 2014).

### RNA extraction and transcript analysis of *OsHAK18*

Isolation of total RNA was extracted from roots, leaf sheaths and blades of treated seedlings with RNAiso plus reagent (TaKaRa, TaKaRa Bio Inc., Dalian, China) following the manufacturer’s instructions. Relative quantitative results were normalized to *OsACT* (accession number AB047313) and are presented here as mean normalized transcript levels obtained by the comparative cycle threshold method (2^−ΔΔCt^). Real-time PCR assays were performed in 20 μL reaction volumes using a QuantStudio 6 flex Real-time PCR system (Applied Biosystems). The primers used are listed in Supplemental Table S1.

### GUS staining and immunostaining analysis

The 2000-bp DNA sequence upstream of the start codon of *OsHAK18* was amplified by PCR and the primers used are listed in Supplemental Table S2. The purified PCR product was then fused upstream of the *GUSPlus* reporter gene in pCAMBIA1300-GN vector via BamHI / KpnI (Chang et al., 2019). The construct was transformed into Agrobacterium strain EHA105 by electroporation, and then transformed into callus derived from mature embryos of japonica rice cultivar Nipponbare via Agrobacterium tumefaciens-mediated transformation (Jia et al., 2011). The GUS staining analysis was performed as described in Ai et al. (2009). Immunostaining analysis with an antibody against GUS was performed as described in Yamaji and Ma (2007), and sections were observed by a confocal laser scanning microscope (Leica, Wetzlar, Germany; TCS SP8X). The cell wall autofluorescence was excited by ultraviolet light with an intensity of 0.7%–3%; the collection bandwidth and gain value were 410–470 nm and 30%–60%, respectively. The collection bandwidth for Alexa Fluor 555 (the secondary antibody) was 560–580 nm, and the gain value was 50%–60% (Dai et al., 2022).

### Subcellular localization assay

For constructing *eGFP*::*OsHAK18* and *OsHAK18*::*eGFP*, the *OsHAK18* coding sequence (including the stop codon) was amplified by PCR using primers listed in Supplemental Table S3. The amplified cDNA was cloned into the middle vector pSAT6A-EGFP-C1 and pSAT6A-EGFP-N1. The middle vector was cut by the PI-PspI and inserted in-frame into the expression vector pRCS2-ocs-nptII. OsRac1-mCherry was used as a plasma membrane control. Transient expression of *OsHAK18* in rice protoplasts, and fluorescence microscopy imaging observation using sequential scanning mode of confocal microscopy (TCS SP8X, Leica, Germany). Excitation/emission wavelengths were 488 nm/498–540 nm for eGFP and 552 nm/600–640nm for mCherry (Dai et al., 2022).

### Yeast strains and culture conditions

Coding sequences of *OsHAK18 and OsHAK5* were cloned into each of the expression vectors p416GPD, pDR196 and pYES2, which contain the constitutive GPD promoter, the inducible PMA promoter and the inducible GAL1 promoter, respectively, for selection and expression in yeast (*Saccharomyces cerevisiae*). The primers used are listed in Supplemental Table S4. The expression vectors were transformed into the K^+^ uptake-deficient R5421 (*MATα, Δtrk1, trk2* :: pCK64, *his3, leu2, ura3, trp1* and *ade2*) strain (Anderson et al. 1992) and the Na^+^-hypersensitive G19 (*MATα*, *his3*, *leu2*, *ura3*, *trp1*, *ade2* and Δena1::HIS3::Δena4) strain (Quintero et al., 1996) of the yeast using the LiAC/ssDNA/PEG method. The transformants were selected on SC-agar plates without uracil (SC-U). Then an AP (arginine and phosphoric acid) medium was used for subsequent growth assays, which were performed as described previously (Horie et al. 2011). For liquid culture of G19, we collected cells at 12 h in 5 mM RbCl and 24 h in 400 mM KCl or NaCl SC-U media for Rb^+^, K^+^ and Na^+^ measurement (Feng et al., 2019).

### Identification of *oshak18* mutant lines and generation of *OsHAK18*-overexpressing transgenic rice

Two different *Tos17* insertion lines of *OsHAK18* knockout mutants in the genetic backgrounds of cv. Nipponbare and cv. Zhonghua11 were requested from NIAS institute, Japan and Huazhong Agricultural University, China, respectively. According to the insertional information, two primers for *OsHAK18* gene and primers specifically for *Tos17* vector were used to determine the *Tos17* insertional sites. Homozygous plants for *Tos17* insertions were identified by two-round RT-PCR analysis (Supplemental Table S5). Further detection of the expression levels of *OsHAK18* in the homozygous mutant seedlings were performed by RT-PCR analysis.

For generating the transgenic rice lines with overexpression of the *OsHAK18* gene, the endogenous promoter of *OsHAK18* (1958 bp sequence upstream of transcriptional start site) was used to replace the ubiquitin promoter in the expression vector pTCK303 (Ai et al., 2009), and the primers used are listed in Supplemental Table S6. Vector harboring *pOsHAK18:OsHAK18* was transferred into *A. tumefaciens* strain EHA105 by electroporation, and the agrobacteria were used to infect callus induced from WT rice (cv. Nipponbare) as described in Tang et al. (2012). A total of 12 *OsHAK18*-OX transgenic lines were selected after GUS staining and PCR analysis by following the protocol described by Yang et al. (2014). Three representative lines (OX1, OX2, OX3) were used for the detailed analysis (Supplemental Figure S4, E and F).

### Measurement of K^+^ and Na^+^ concentrations in xylem sap and phloem sap

Similar-size 10-day-old rice seedings of each WT and transgenic lines were grown in the IRRI solution for 8 weeks and then transferred to 1- or 0.1-mM K^+^ solutions for 3 days before the sampling. The detailed protocols for the shoot excision, the xylem sap collection by cotton ball, and the determination of K^+^ concentration were described previously by Yang et al. (2014). At the same time, the shoot part removed from plants during the collection of the xylem sap was put into calibrated test tubes containing 20 mL 20 mM EDTA solution (pH=7.5-8.0), and placed in a dark environment with air humidity higher than 70% for 5 hours to collect the phloem sap. The protocol was derived from King and Zeevaart (1974). The amounts of K^+^, Na^+^ and glutamine released in the EDTA solutions were determined by ICP-OES (Agilent 710, Agilent Technologies, USA) and Agilent 1200 HPLC coupled with Agilent G6410 mass spectra, respectively. The concentration ratio of K^+^ or Na^+^ to glutamine was used to standardize the K^+^ or Na^+^ data (Ren et al., 2005).

### Detecting the effect of *OsHAK18* expression level on the rice root short-term K^+^ acquisition

Rice K^+^ depletion experiments followed the previously described procedures (Nieves-Cordones et al., 2007; Rubio et al., 2008) with some modifications. Similar-size 10-day-old rice seedlings were grown in the IRRI solution containing 0.1- or 1-mM K^+^ for 4 weeks, and then subjected to K^+^ starvation for 7 days by excluding K^+^ from both solutions. After 7 days of K^+^ starvation, the roots were rinsed with a cold K^+^-free nutrient solution, and then seedlings of WTs or transgenic lines were transferred to containers filled with 350 mL of solution for 0.1- or 1-mM K^+^ treatment (two plants per container). Three mL of the solution were collected from each container at 0, 0.5, 1, 2, 4, 6, 8 and 12 h and K^+^ concentration in the sample was determined by ICP-OES (Agilent 710, Agilent Technologies, USA) as described previously (Cai et al., 2012). At the end of the experiment, root fresh weight of each plant was determined. K^+^ acquisition rate was calculated by taking the amount of K^+^ decrease in the solution at every time point and normalizing it to the root fresh weight (g) of the corresponding individual plant (Chen et al., 2015).

### Split-root experiment for assay of Rb^+^ circulation

Similar-size 10-day-old rice seedings of each WT and transgenic lines were grown in the IRRI solution for 6 weeks. Then, the rice plants were grown individually in pots, and the roots per plant were divided almost equally into two plastic vessels containing a nutrient solution of 0.5 mM Rb^+^ (Root-Rb^+^) and 0.5 mM K^+^ (Root-K^+^), respectively. After 3 hours, both roots were placed back into 0.5 mM K^+^ nutrient solution for another 3 hours. The two root parts and the shoots were collected at 3 h and 6 h, counting from the beginning of Rb^+^ treatment and their Rb^+^ content was measured by ICP-MS (Thermo Fisher Scientific, USA).

### Measurement of soluble sugar content

Roots, stems, leaf sheaths and blades of soil-grown plants were harvested at the mature stage. All samples were snap-frozen in liquid nitrogen and stored at −80°C. Soluble sugar was extracted with 80% (v/v) ethanol from an 80°C water bath. Soluble content was determined based on the method described by Leyva et al. (2008) with minor modifications using anthrone-sulfuric acid colorimetry. The anthrone reagent was prepared right before analysis by dissolving 0.1 g of anthrone (0.1%) in 100 mL of 70% (v/v) sulfuric acid, protected from light and used within 12 h. The reaction liquid was heated and then cooled to room temperature before reading absorbance at 625 nm.

### Statistical analysis

Data were analyzed by non-paired, two-tailed, equal-variance t-test using the SPSS 21 program (SPSS Inc., Chicago, IL, USA).

## Acknowledgements

This work was supported by National Key Research and Development Program of China (grant no. 2021YFF1000402-1) and National Natural Science Foundation of China (grant no. 31872166).

## Data availability

The data that support the findings of this study are available from the corresponding authors upon reasonable request.

## Author contributions

L.Y. and G.X. conceived the study and designed the experiments; L.P. conducted most of the experiments; H.X., R.L. and Y.Z. generated transgenic lines and performed part of the experiments; M.G. conducted the immunostaining assay; L.Y., G.X. and L.P. analyzed the data and together with N.M. wrote the manuscript.

## Supplemental data

Supplemental data associated with this article can be found in the online version.

**Supplemental Figure S1.** Phylogenetic tree of HAK/KUP/KT family genes in rice (OsHAK) and Arabidopsis.

**Supplemental Figure S2.**. *OsHAK18* expression and tissue localization in rice.

**Supplemental Figure S3.** Complementation of a yeast mutant R5421 defective in K^+^ uptake by *OsHAK18*.

**Supplemental Figure S4.** Homozygous *Tos17* insertion mutants of *OsHAK18* gene in rice and characterization of *OsHAK18*-overexpressing (OX) transgenic rice lines in the background of the Nipponbare cultivar.

**Supplemental Figure S5.** Effect of *OsHAK18* expression level on rice growth at seedling stage.

**Supplemental Figure S6.** Effect of *OsHAK18* expression level on total K^+^ content at the seedling stage.

**Supplemental Figure S7.** Effect of *OsHAK18* expression level on short-term root K^+^ acquisition rates at different K^+^ supply conditions.

**Supplemental Figure S8.** Effect of *OsHAK18* expression level on agronomic traits at reproduction.

**Supplemental Figure S9.** Effect of *OsHAK18* expression level on salt sensitivity of rice.

**Supplemental Figure S10.** Effect of *OsHAK18* expression level on K^+^ and Na^+^ content in rice supplied with NaCl in the culture solution.

**Supplemental Table S1.** Primers used to amplify *OsACT* and *OsHAK18* for quantitative real-time PCR.

**Supplemental Table S2.** Primers used to amplify *OsHAK18* promoter for tissue localization.

**Supplemental Table S3.** Primers used to amplify *OsHAK18* cDNA for subcellular localization.

**Supplemental Table S4.** Primers used to amplify *OsHAK18* and *OsHAK5* cDNA for functional assay in yeast.

**Supplemental Table S5.** Primers for identification of two homozygous mutant lines of *oshak18*.

**Supplemental Table S6.** Primers used to amplify *OsHAK18* promoter and cDNA for construction of over-expression.

