## Supplemental Figures for "Potassium transporter OsHAK18 mediates shoot-to-root circulation of potassium and sodium and source-to-sink translocation of soluble sugar in rice"

### Supplemental Figures and Tables

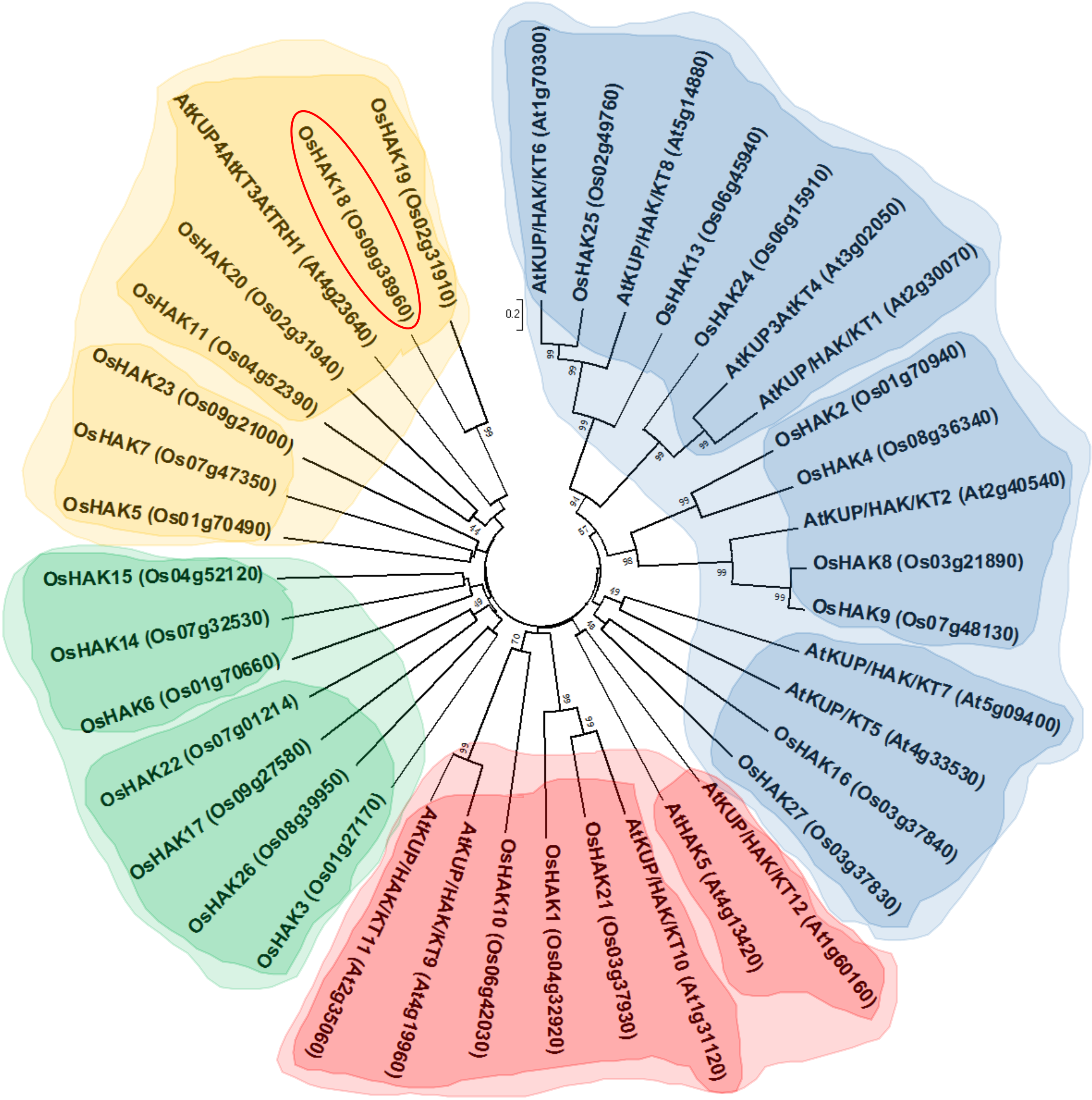


**Figure S1. Phylogenetic tree of HAK/KUP/KT family genes in rice (OsHAK) and Arabidopsis.** The joint unrooted tree was generated using MEGA7 by the neighborjoining method.

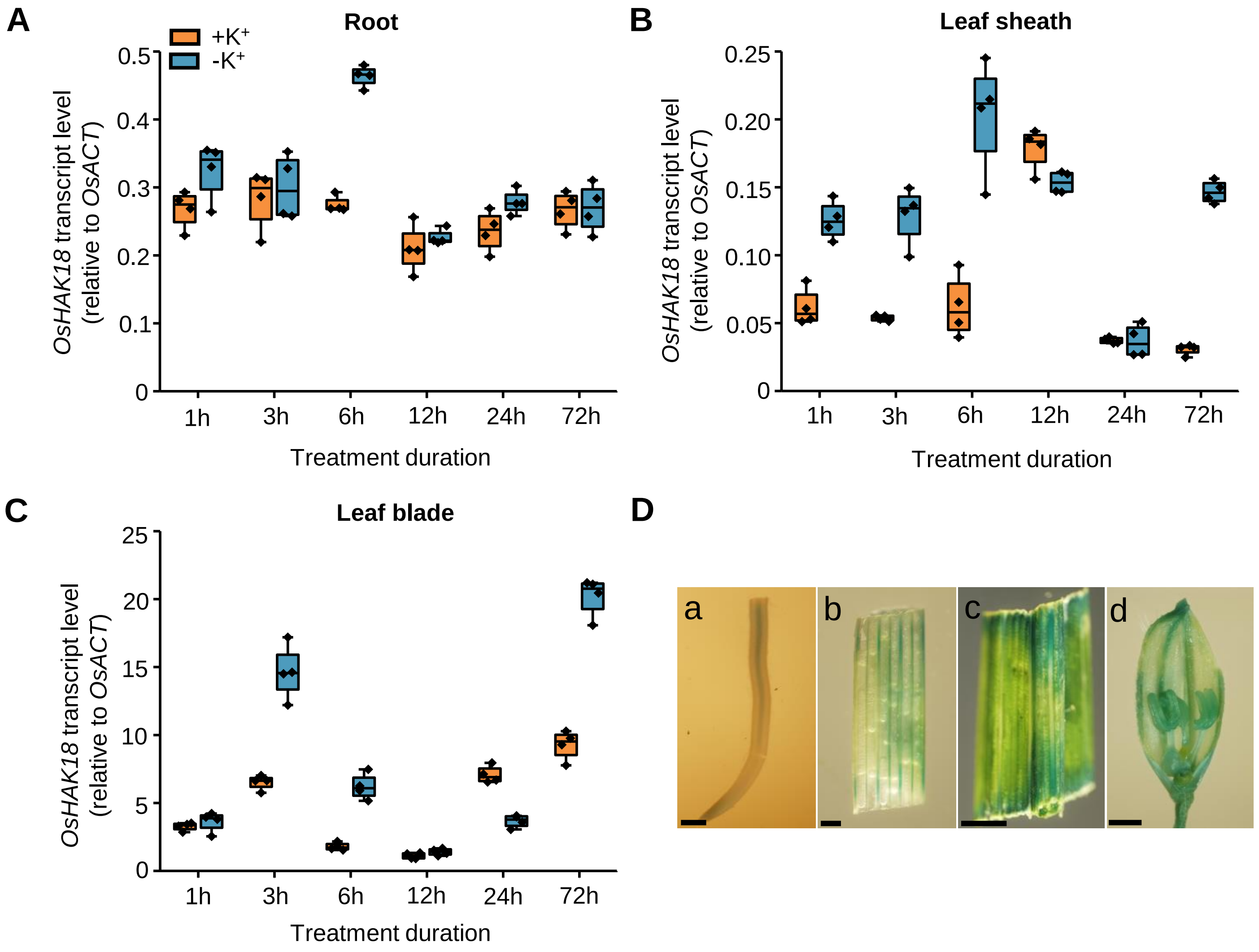


**Figure S2. *OsHAK18* expression and tissue localization in rice.**
A-C, Organ-specific expression analysis of *OsHAK18* by reverse transcriptase-quantitative polymerase chain reaction (qRT-RCR) using RNA from 3-week-old seedlings of japonica cultivar Nipponbare treated with $+K^+$ (1 mM $K^+$) or $-K^+$ (0 mM $K^+$) nutrient solution for the indicated durations. *OsHAK18* expression in roots (A), leaf sheaths (B) and leaf blades (C) was analyzed. In the box plots, the symbols represent individual data (biological repeats), whiskers represent minimum and maximum data values, the boxes are delimited by the first and the third quartiles, and the horizontal line within the box indicates the median. D, Tissue-specific localization of *OsHAK18* as indicated by the GUS reporter gene in transgenic rice plants harboring the *ProOsHAK18:GUS* construct. a, root; b, leaf sheath; c, leaf blade; d, glumous flower. Scale bars = 1 mm.

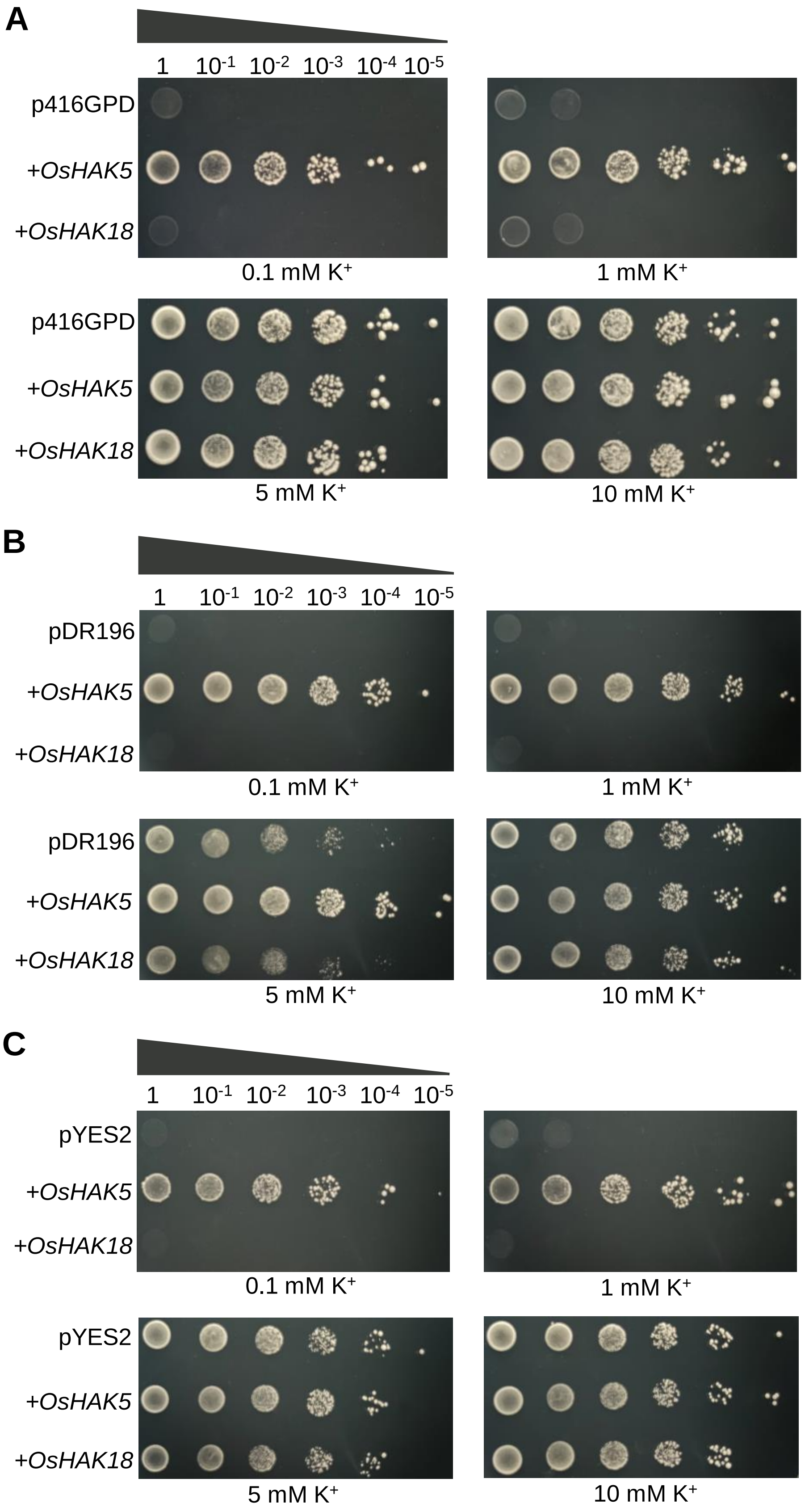


**Figure S3. Complementation of a yeast mutant R5421 defective in $K^+$ uptake by *OsHAK18*.**
Yeast cells harboring either an *OsHAK5* construct (positive control) or an *OsHAK18* construct in one of the vectors p416GPD (A), pDR196 (B) and pYES2 (C), or the corresponding empty vector, were grown in synthetic drop-out (-Ura) liquid medium containing 2% (w/v) glucose or galactose to an optical density of 1 at the wavelength of 600 nm (OD600). Then 5-uL aliquots of 10-fold serial dilutions were spotted on agar plates containing different $K^+$ concentrations. Plates were incubated at 30℃ for 4 d.

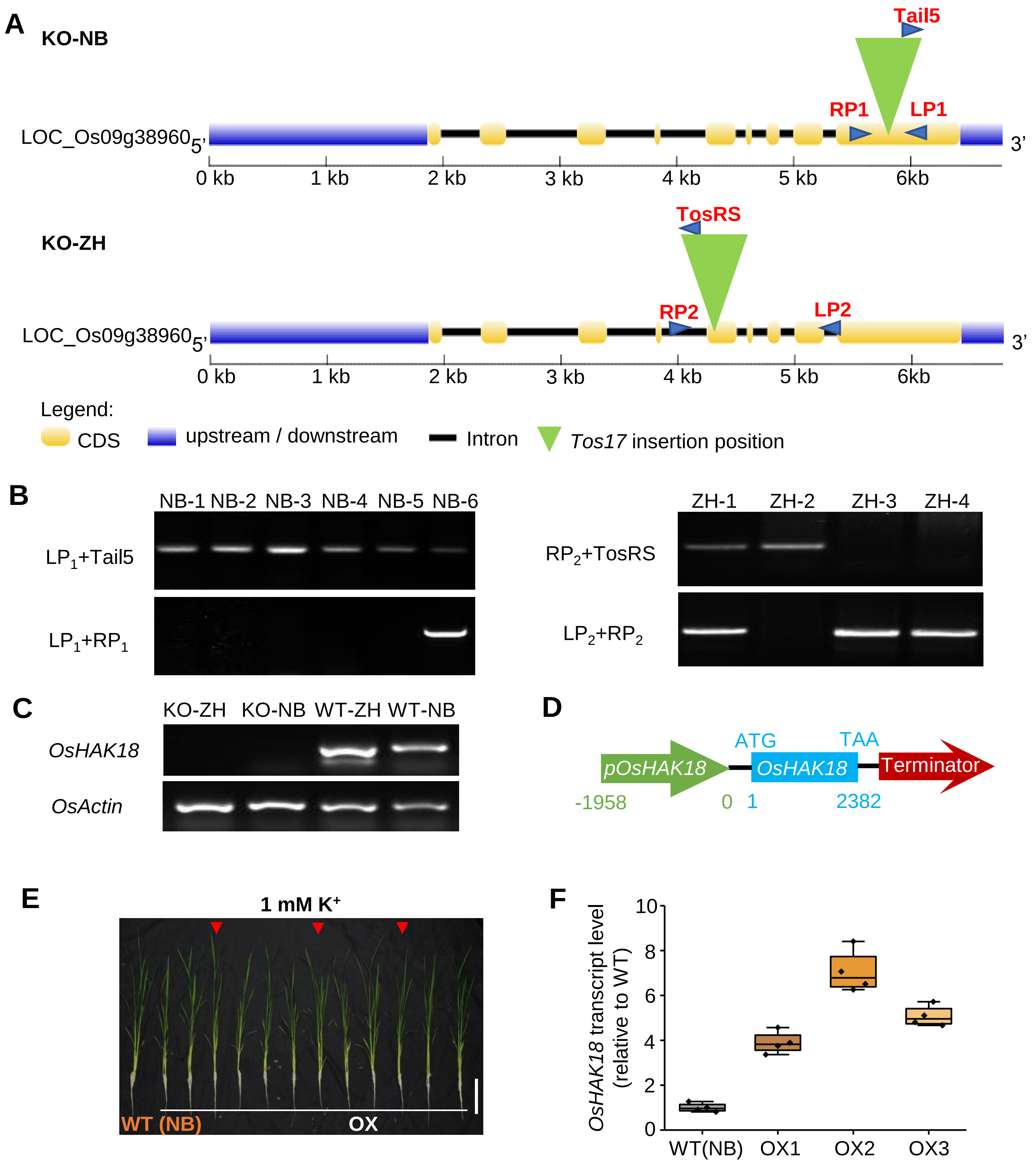


**Figure S4. Homozygous *Tos17* insertion mutants of *OsHAK18* gene in rice and characterization of *OsHAK18*-overexpressing (OX) transgenic rice lines in the background of the Nipponbare cultivar.**

A, A schematic diagram shows the *Tos17* insertion positions of *oshak18* mutants in genetic background of cv. Nipponbare (NB) and cv. Zhonghua11 (ZH). B, Verification of plants homozygosity (homo) for each of the *Tos17* inserts using two rounds of RT-PCR. C, Expression levels of *OsHAK18* in leaves of Nipponbare (NB) and Zhonghua11 (ZH) WTs and *oshak18* mutants (KO). RT-PCR was performed on total extracted RNA. D, Construction of *pOsHAK18::OsHAK18* expression vector. E, Growth performance of *OsHAK18*-overexpressing transgenic rice lines (under self-promoter) and WT plants for 6 weeks. Scale bars = 10 cm. F, Relative transcript level of *OsHAK18* in shoots of OX lines and WT plants using total RNA isolated from the leaves of two weeks-old seedlings. The box plots details are as in Figure 2.

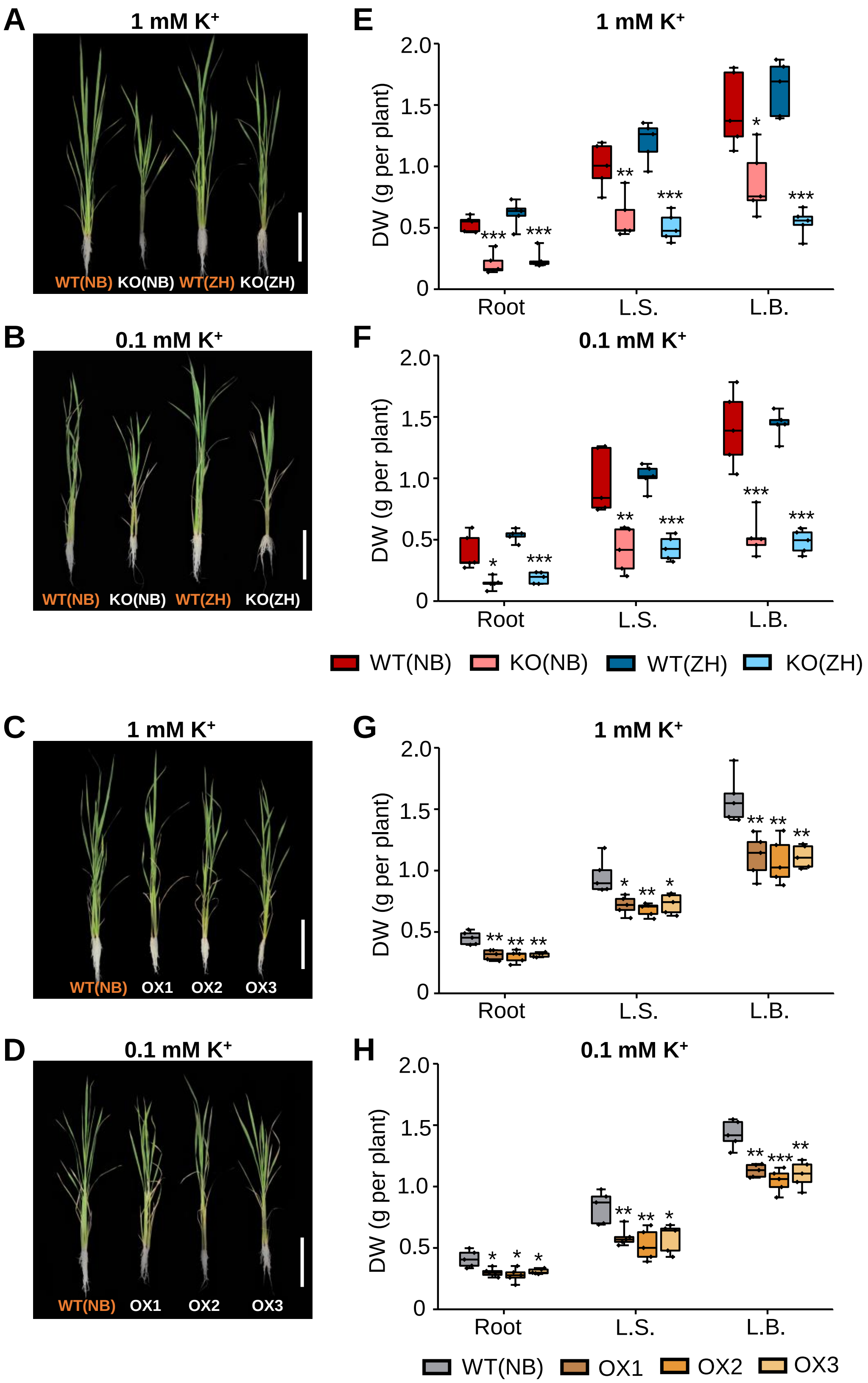


**Figure S5. Effect of *OsHAK18* expression level on rice growth at seedling stage.**
Ten-day-old seedlings were grown in IRRI solution containing 1 mM $K^+$ for 4 weeks, then transferred to 1 mM $K^+$ (A, C, E, G) or to 0.1 mM $K^+$ (B, D, F, H) solutions for 2 weeks. .A-D, Growth performance. Scale bars = 10 cm. E-H, Dry weight. The box plots and t-tests are as in Figure 2. Significant differences from the corresponding wild type (WT) are indicated by asterisks (*, $P < 0.05$; **, $P < 0.01$; ***, $P < 0.001$).

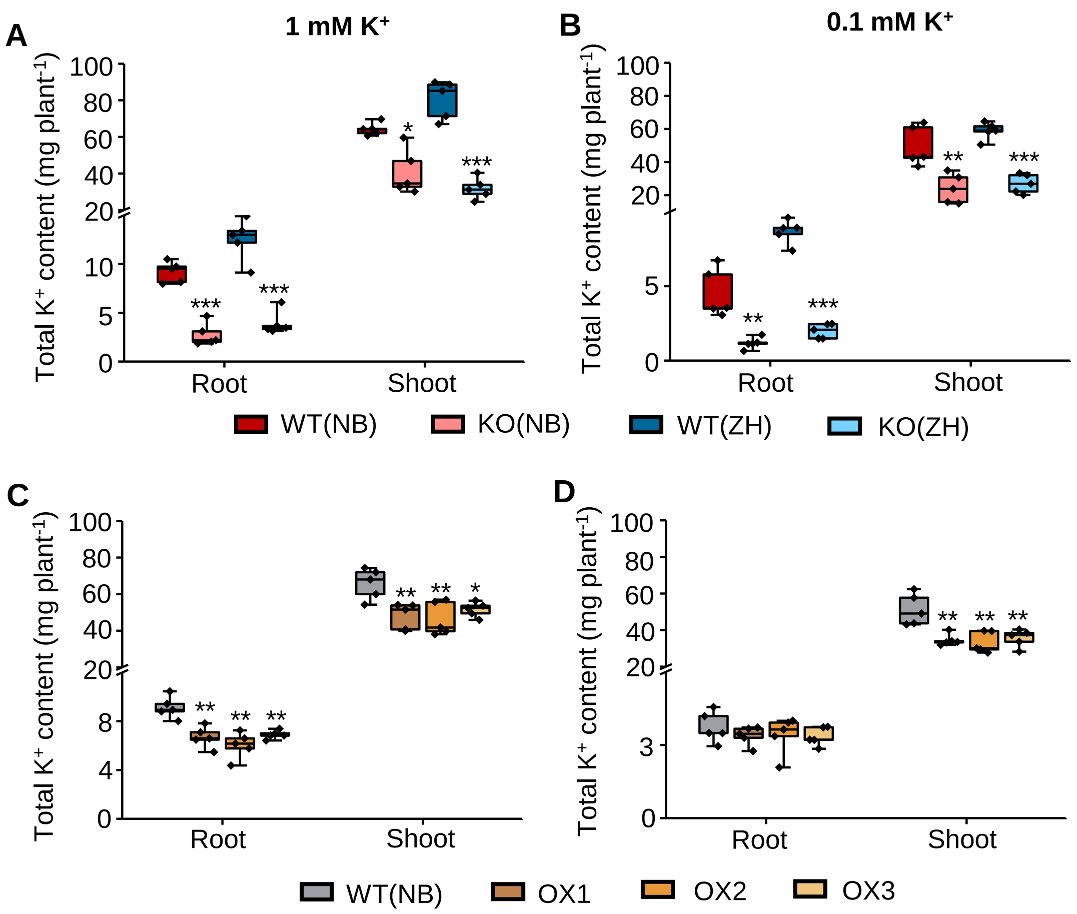


**Figure S6. Effect of *OsHAK18* expression level on total K⁺ content at the seedling stage.** Ten-day-old seedlings were grown in IRRI solution containing 1 mM $K^+$ for 4 weeks, then transferred to 1 mM $K^+$ (A, C) or 0.1 mM $K^+$ (B, D) solutions for 2 weeks. A-D, Total $K^+$ content in root and shoot. The box plots and t-test details are as in Figure 2. Significant differences from the corresponding WT are indicated by asterisks (*, $P < 0.05$; **, $P < 0.01$; ***, $P < 0.001$).

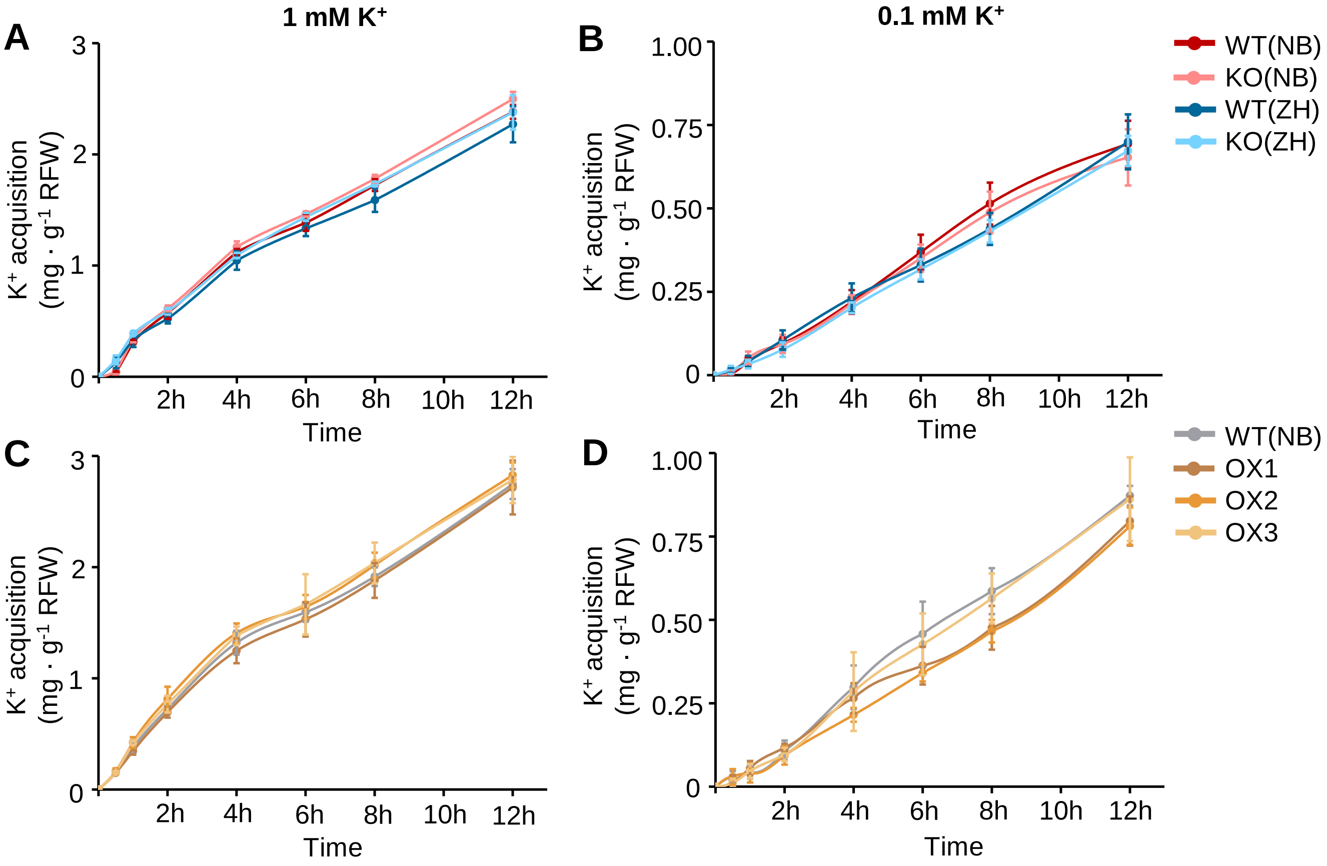


**Figure S7. Effect of *OsHAK18* expression level on short-term root $K^+$ acquisition rates at different $K^+$ supply conditions.**

Ten-day-old rice seedlings were grown in the IRRI solution with 1 or 0.1 mM $K^+$ for 4 weeks and then subjected to $K^+$ starvation for 7 days before conducting a $K^+$ acquisition test. A and B, KO lines and their WT. C and D, Overexpression lines and their WT. $K^+$ concentration in the culture solution with initial $K^+$ at 1 mM (A, C) and 0.1 mM (B, D), respectively. RFW, Root fresh weight. Error bars indicate SD (n = 4).

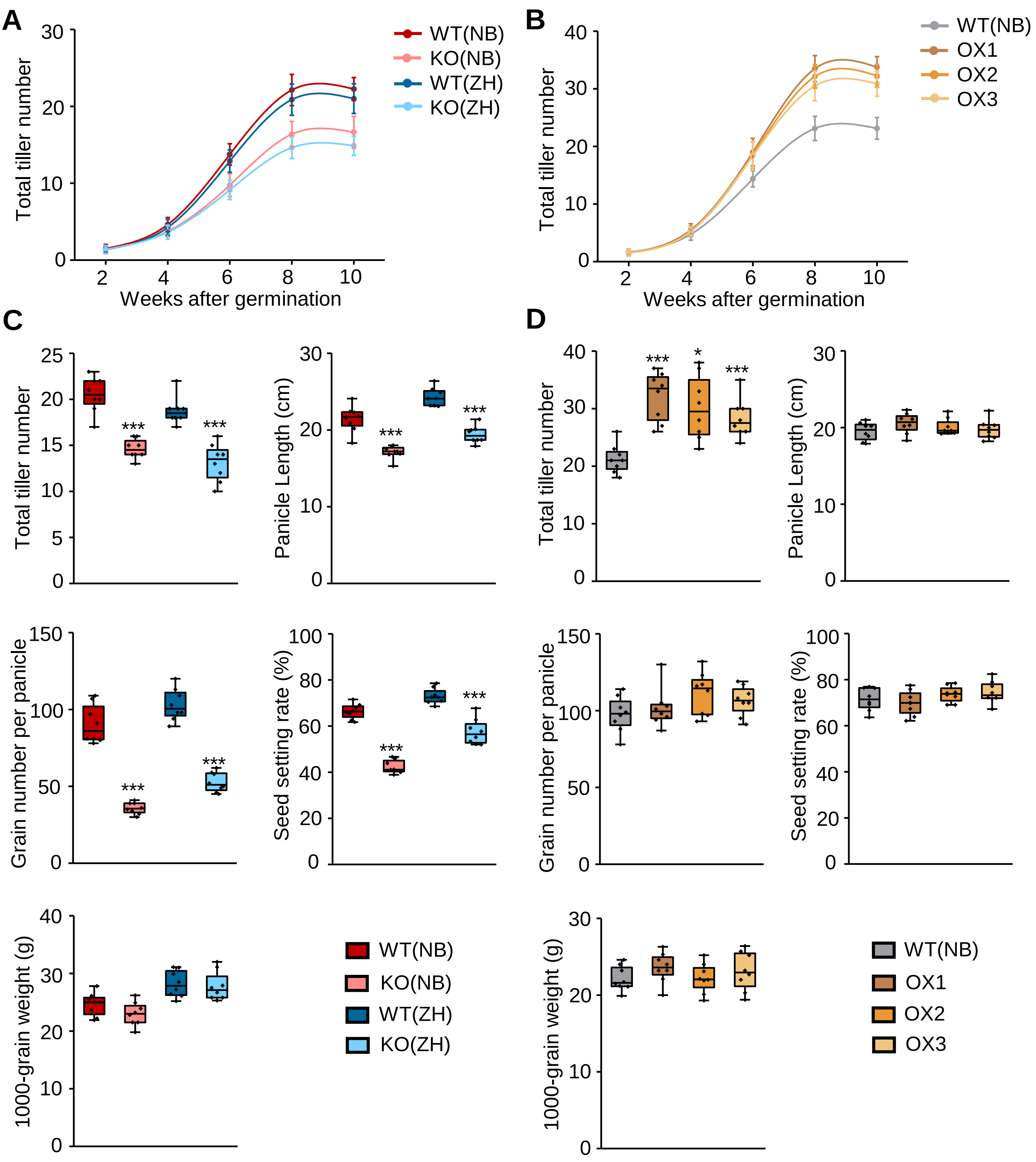


**Figure S8. Effect of *OsHAK18* expression level on agronomic traits at reproduction.**
Rice plants were grown in the paddy field and irrigated, fertilized and medicated regularly until they were fully mature (approx. 20-weeks-old). A and B, Time course of the total tiller number increase during about 10 weeks of growth after seed germination within each cultivar (compared separately at each time point). Error bars indicate SD (n = 8). C and D, Total tiller number, panicle length, seed setting rate, grain number per plant and 1000-grain weight. The box plots and t-test details are as in Figure 2. Significant differences from the corresponding WT are indicated by asterisks (*, $P < 0.05$; **, $P < 0.01$; ***, $P < 0.001$).

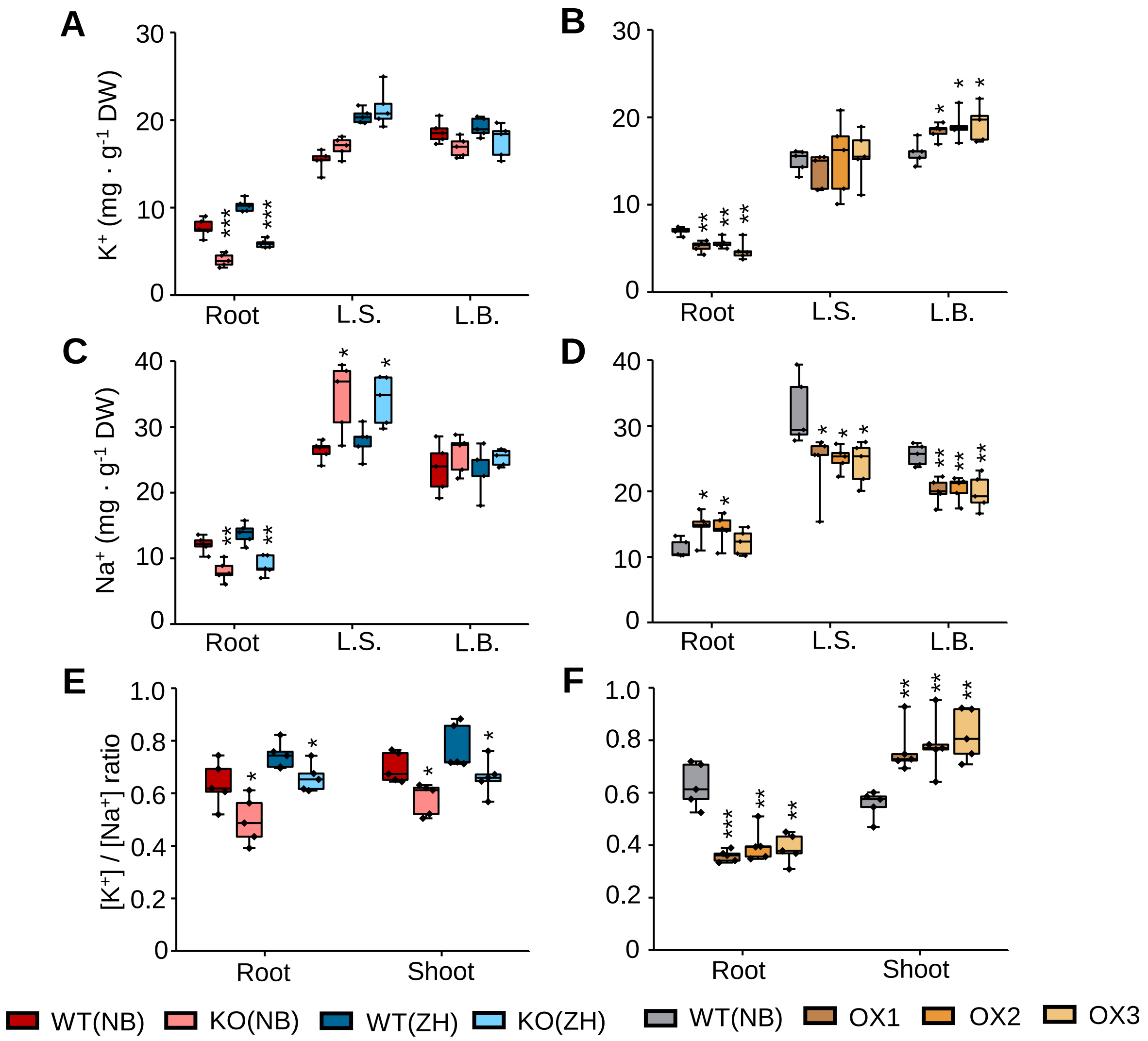


**Figure S9. Effect of *OsHAK18* expression level on salt sensitivity of rice.**
Ten-day-old seedlings were grown in IRRI solution containing 1 mM $K^+$ for 4 weeks, then transferred to a solution containing additionally 100 mM NaCl and allowed to grow for 2 weeks. L.S., Leaf sheath; L.B., Leaf blade. The box plots and t-test details are as in Figure 2. Significant differences from the corresponding WT are indicated by asterisks (*, $P < 0.05$; **, $P < 0.01$; ***, $P < 0.001$).

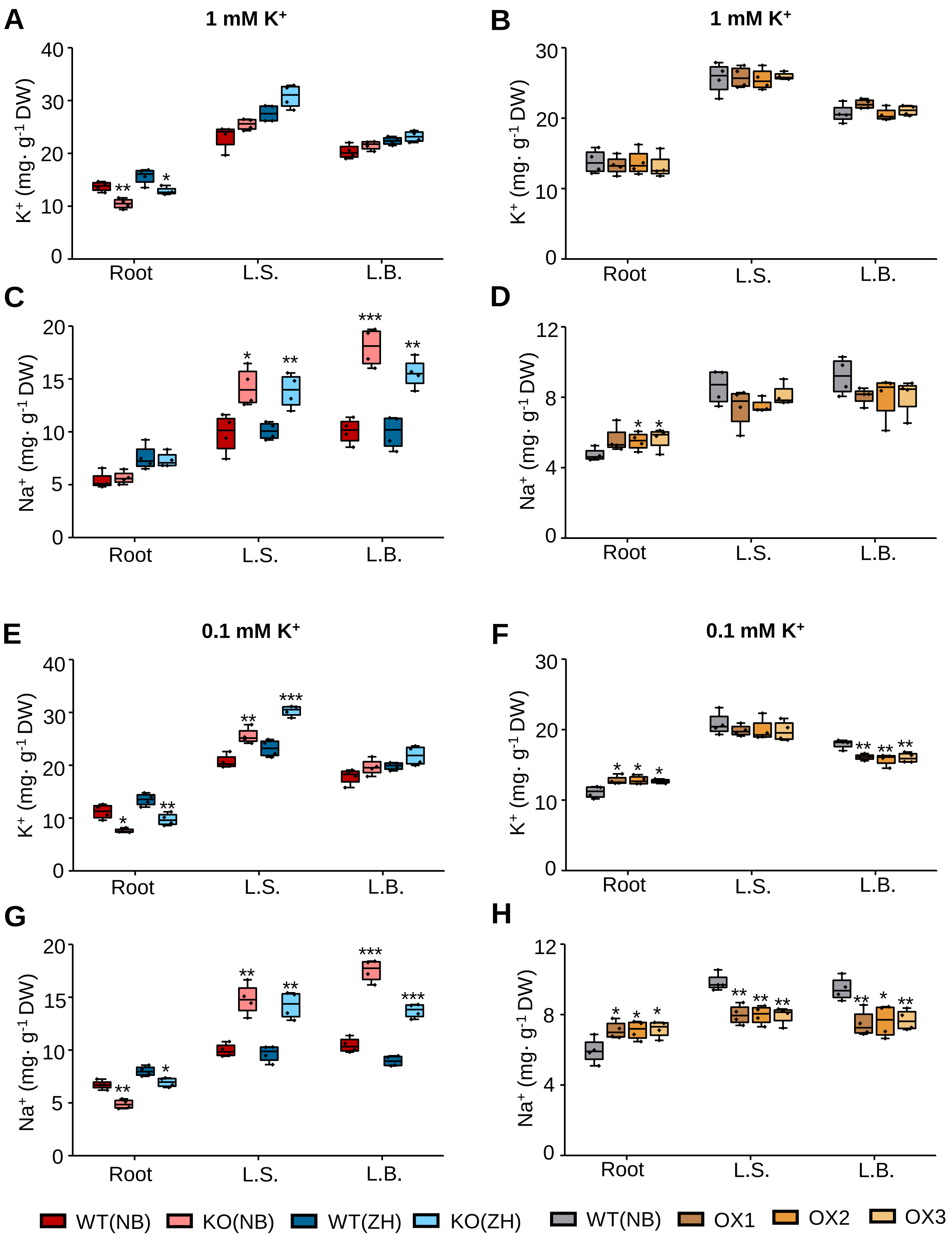


**Figure S10. Effect of *OsHAK18* expression level on $K^+$ and $Na^+$ content in rice supplied with NaCl in the culture solution.**

Ten-day-old seedlings were grown in IRRI solution containing 1 mM $K^+$ for 2 weeks, then transferred to 1 mM $K^+$ (A-D) or to 0.1 mM $K^+$ (E-H) solutions for 2 weeks, then transferred to 1 mM $K^+$ or to 0.1 mM $K^+$ solutions containing additionally 20 mM NaCl and allowed to grow for 3 d, then returned to 1 mM or 0.1 mM $K^+$ medium without NaCl for additional 3 d. L.S., Leaf sheath; L.B., Leaf blade. The box plots and t-tests are as in Figure 2. Significant differences from the corresponding WT are indicated by asterisks (*, $P < 0.05$; **, $P < 0.01$; ***, $P < 0.001$).

**Table S1. Primers used to amplify *OsACT* and *OsHAK18* for quantitative real-time PCR.**

| Gene (Accession No.) | Primer sequences (5' to 3') |
|---|---|
| *OsACT* (AB047313) | F: TTATGGTTGGGATGGGACA |
| | R: AGCACGGCTTGAATAGCG |
| *OsHAK18* (AK065464) | F: CTCGCACTTCATCACCAA |
| | R: TTCACCAGGAACCTCTCAT |

**Table S2. Primers used to amplify *OsHAK18* promoter for tissue localization.**

| Target vector | Primer sequences (5' to 3') |
|---|---|
| pCAMBIA1300-GN | F:GCCTGCAGGTCGACTCTAGAGGATCCTGTCCCACAGATCTTATTGT |
| | R: TTTACCCTCAGATCTACCATGGTACCGGGTTCAGACTTCAGATCAA |

Notes: The incorporated two restriction sites sequences of BamHI (GGATCC) and KpnI (GGTACC) are underlined.

**Table S3. Primers used to amplify *OsHAK18* cDNA for subcellular localization.**

| Target vector | Primer sequences (5' to 3') |
|---|---|
| pSAT6A-EGFP-C1 | F: AAGCTTCTATGGAGACCAGAACAAATGAGTAT |
| | R: CCCGGGCTACACGTAGAAAACCTGCCCA |
| pSAT6A-EGFP-N1 | F: AAGCTTATGGAGACCAGAACAAATGAGTAT |
| | R: CCCGGGCACGTAGAAAACCTGCCCA |

Notes: The incorporated two restriction sites sequences of HindIII (AAGCTT) and SmaI (CCCGGG) are underlined.

**Table S4. Primers used to amplify *OsHAK18* and *OsHAK5* cDNA for functional assay in yeast.**

| Product | Primer sequences (5' to 3') |
|---|---|
| *OsHAK18*-cDNA | F-H: AAGCTTAACACAATGTCTATGGAGACCAGAACAAATGAGTATT |
| | F-X: TCCCCCCGGGAACACAATGTCTATGGAGACCAGAACAAATGAGTATT |
| | R: TACGAATTCTTACACGTAGAAAACCTGCCCAACA |
| *OsHAK5*-cDNA | F-H: ATCAAGCTTAACACAATGTCTATGACCGAGCCTCTGCACACAA |
| | F-X: TCCCCCCGGGAACACAATGTCTATGACCGAGCCTCTGCACACAA |
| | R: TACGAATTCCTAGATCTCGTACGTCATTCCT |

Notes: The incorporated two restriction sites sequences of HindIII (AAGCTT), XmaI (CCCGGG), and EcoRI (GAATTC) are underlined.

**Table S5. Primers for identification two homozygous mutant lines of *oshak18*.**

| Gene (Accession No.) | Genetic background | Primer sequences (5' to 3') |
|---|---|---|
| *OsHAK18* (AK065464) | cv. Nipponbare | LP1: ATCCCTGGGTTATTTGGGAG |
| | | RP1: GCTGTCATTGCTTGTGGAGA |
| | | Tail-5: CATCGGATGTCCAGTCCATTG |
| | cv. Zhonghua11 | LP2: AAAATCAGCCAGGCTTGAAG |
| | | RP2: TGCAAACAAGTAATGCGGAG |
| | | TosR5: GAAGGGGGGTGTTAAATATATATAC |

**Table S6. Primers used to amplify *OsHAK18* promoter and cDNA for construction of over-expression.**

| Product | Primer sequences (5' to 3') |
|---|---|
| *OsHAK18*-promoter | F: GTAAAACGACGGCCAGTGCCAAGCTTTGTCCCACAGATCTTATTGT |
| | R: TCATTTGTTCTGGTCTCCATGGTACCGGGTTCAGACTTCAGATCAA |
| *OsHAK18*-cDNA | F: TCGACTCTAGAGGATCCCCGGGTACCATGGAGACCAGAACAAATGA |
| | R: TCATGGTCTTTGTAGTCCATACTAGTCACGTAGAAAACCTGCCCAA |

Notes: The incorporated three restriction sites sequences of HindIII (AAGCTT), KpnI (GGTACC) and SpeI (ACTAGT) are underlined.
